# Open science resources from the Tara Pacific expedition across coral reef and surface ocean ecosystems

**DOI:** 10.1101/2022.05.25.493210

**Authors:** Fabien Lombard, Guillaume Bourdin, Stéphane Pesant, Sylvain Agostini, Alberto Baudena, Emilie Boissin, Nicolas Cassar, Megan Clampitt, Pascal Conan, Ophélie Da Silva, Céline Dimier, Eric Douville, Amanda Elineau, Jonathan Fin, J. Michel Flores, Jean François Ghiglione, Benjamin C.C. Hume, Laetitia Jalabert, Seth G. John, Rachel L. Kelly, Ilan Koren, Yajuan Lin, Dominique Marie, Ryan McMinds, Zoé Mériguet, Nicolas Metzl, David A. Paz-García, Maria Luiza Pedrotti, Julie Poulain, Mireille Pujo-Pay, Joséphine Ras, Gilles Reverdin, Sarah Romac, Alice Rouan, Eric Röttinger, Assaf Vardi, Christian R. Voolstra, Clémentine Moulin, Guillaume Iwankow, Bernard Banaigs, Chris Bowler, Colomban de Vargas, Didier Forcioli, Paola Furla, Pierre E. Galand, Eric Gilson, Stéphanie Reynaud, Shinichi Sunagawa, Matthew B. Sullivan, Olivier Thomas, Romain Troublé, Rebecca Vega Thurber, Patrick Wincker, Didier Zoccola, Denis Allemand, Serge Planes, Emmanuel Boss, Gaby Gorsky

## Abstract

The Tara Pacific expedition (2016-2018) sampled coral ecosystems around 32 islands in the Pacific Ocean and the ocean surface waters at 249 locations, resulting in the collection of nearly 58,000 samples. The expedition was designed to systematically study warm coral reefs and included the collection of corals, fish, plankton, and seawater samples for advanced biogeochemical, molecular, and imaging analysis. Here we provide a complete description of the sampling methodology, and we explain how to explore and access the different datasets generated by the expedition. Environmental context data were obtained from taxonomic registries, gazetteers, almanacs, climatologies, operational biogeochemical models, and satellite observations. The quality of the different environmental measures has been validated not only by various quality control steps but also through a global analysis allowing the comparison with known environmental large-scale structures. Such a wide released datasets opens the perspective to address a wide range of scientific questions.

## Background & Summary

Marine ecosystems are facing numerous perturbations either of seasonal, climatic, or biological origin which are now overamplified by perturbations due to anthropogenic activities. The resilience of marine ecosystems to perturbations is a general concern, especially when providing ecosystem services and supporting human activities. Tropical coral reefs maintain important ecological services such as fisheries, tourism, or coastal protection, but are also among the most sensitive ecosystems to environmental changes^1, 2^. Furthermore, coral health is not only governed by the environment, but also by the holobiont and its symbiotic interactions encompassing a wide range of eukaryotic organisms (e.g., crustaceans, molluscs, fishes), endosymbiotic microalgae, bacteria, fungi, and viruses. In the open sea, coral ecosystems are associated with islands and participate in their long-term ecological and geological resilience. Coral ecosystems are hotspots of biological activities and energy flux that have a strong effect on the open sea through nutrient enrichment that could propagate in the open ocean, supporting fisheries or biogeochemical fluxes in other marine ecosystems^3, 4^.

However, a more complete understanding of how coral ecosystems are reacting to environmental stresses is complicated as multiple spatial (from microscale to mesoscale) and temporal (from minutes, day, seasons or decades) scales are involved, as well as various biological complexity levels (from molecular, genetic, physiological to ecosystem). Monitoring ecosystems features at large biological, spatial, and temporal scales is very challenging. An alternative is to use “space-for-time” substitutions which assumes that processes observed at various static spatial scales could reflect what could happen if the same ecological forcing happens at various temporal scales^5^. Historically, this method was used for centuries, for example when Charles Darwin used it to describe the development of islands from barrier reefs, fringing reefs to atolls^6^. This method is still commonly used in ecology, notably when species distribution^7^ or even diversity^8^ are modelled using niche models.

This type of approach is often limited by the compatibility between datasets, where many observations often originated from separate studies with heterogeneous protocols, methods or measurements. In this respect, large global expeditions have often paved the way to major scientific breakthroughs from the early expeditions conducted by the Beagle or HSM challenger to the more recent Malaspina or Tara Ocean expeditions^9–11^.

The Tara Pacific expedition has applied a pan-ecosystemic approach on coral reefs and their surrounding waters at the entire ocean basin scale throughout the Pacific Ocean^12^. The aim was to propose a baseline reference of coral holobiont genomic, transcriptomic and metabolomic diversity spanning from genes to organisms and its interaction with the environment. Tara Pacific focused on widely distributed organisms, two scleractinian corals (*Pocillopora meandrina* and *Porites lobata*), one hydrocoral (*Millepora platyphylla*) and two reef fish (*Acanthurus triostegus* and *Zanclus cornutus*) together with their contextual biological (plankton) and physicochemical environment^13^.

The collaboration of more than 200 scientists and participants during this expedition, made it possible to sample coral systems across 32 islands (102 sites), together with 249 oceanic stations, resulting in a collection of 57859 samples encompassing the integral study of corals, fishes, plankton, and seawater. As with previous Tara expeditions^14^, organizing and cross-linking the various measurements is a stepping-stone for open-access science resources following FAIR principles (Findable Accessible Interoperable and Reusable^15^). In this effort, the strategy adopted by Tara Pacific is to provide open access data and early and full release of the datasets once validated or published. Such an approach ensures a long-lasting preservation, discovery and exploration of data by the scientific community which will certainly lead to new hypotheses and emerging concepts.

Here we present an overview of the sampling strategy used to collect coral holobiont in connection with its local, large scale or historical environment. We also provide a critical assessment of the environmental context. We provide the full registries describing the geospatial, temporal, and methodological information for every sample, and connect it to the various sampling events or stations. Extensive environmental context is also provided at the level of samples or stations. Such registries and environmental context collections are essential for researchers to explore the Tara Pacific data and will be updated and complemented when additional datasets will be released to the public. Throughout the entire manuscript, terms stated [within brackets] refers to the terms used within the registry or in environmental context datasets.

## Methods

### 1 Sampling locations

Tara Pacific aimed to deploy the same sampling and analysis protocol at large scale to offer a comparative suite of samples covering the widest environmental envelope while optimizing cruising and sampling time over the 2.5 years of the sampling effort. Protocols and global objectives of the Tara Pacific expedition were previously mentioned^12^ for coral samples and are detailed here in connection with the sample registry. Similarly protocols and global objectives for ocean and atmosphere sampling were previously described^13^ for the 249 stations sampled during daytime (noted [OA001] to [OA249]; nighttime sampling between stations and other non-systematic sampling events were noted [OA000]).

A set of 32 island systems (noted [I01] to [I32] in registry; Table 1, Figure 1) were targeted to cover the widest range of conditions as possible, from temperate latitudes to the equator, from the low diversified system of the eastern Pacific to the highly diverse western Pacific warm pool^16^. The variety of coral reef systems explored includes continental islands, remote volcanic islands up to atolls, with varying island sizes or human populations (Table 1). Generally, 3 sites ([S01] to [S03]) per island were selected to conduct the full sampling strategy within 4 days. Occasionally only 2 or up to 5 sites were selected (Table 1).

**Figure 1.**
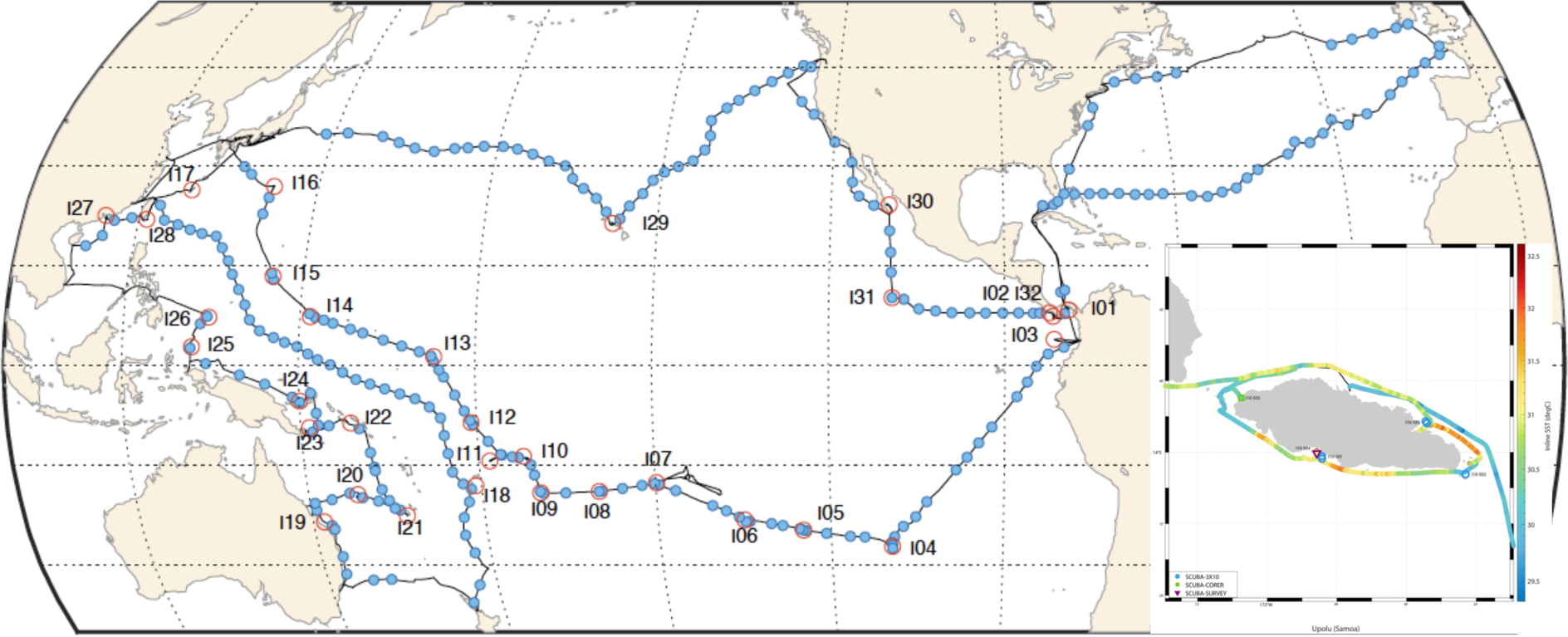
Tara Pacific expedition (2016-2018) sampling map. A) Map of sampled coral systems (red circles) and oceanic stations (blue dots). B) example of coral sampling locations around Upolu (Samoa; I10) with overlaid temperature as recorded by the inline thermosalinograph on the boat trajectory. The absence of sampling in the middle of the return trip in the Atlantic Ocean is due to bad weather.

**Table 1:**
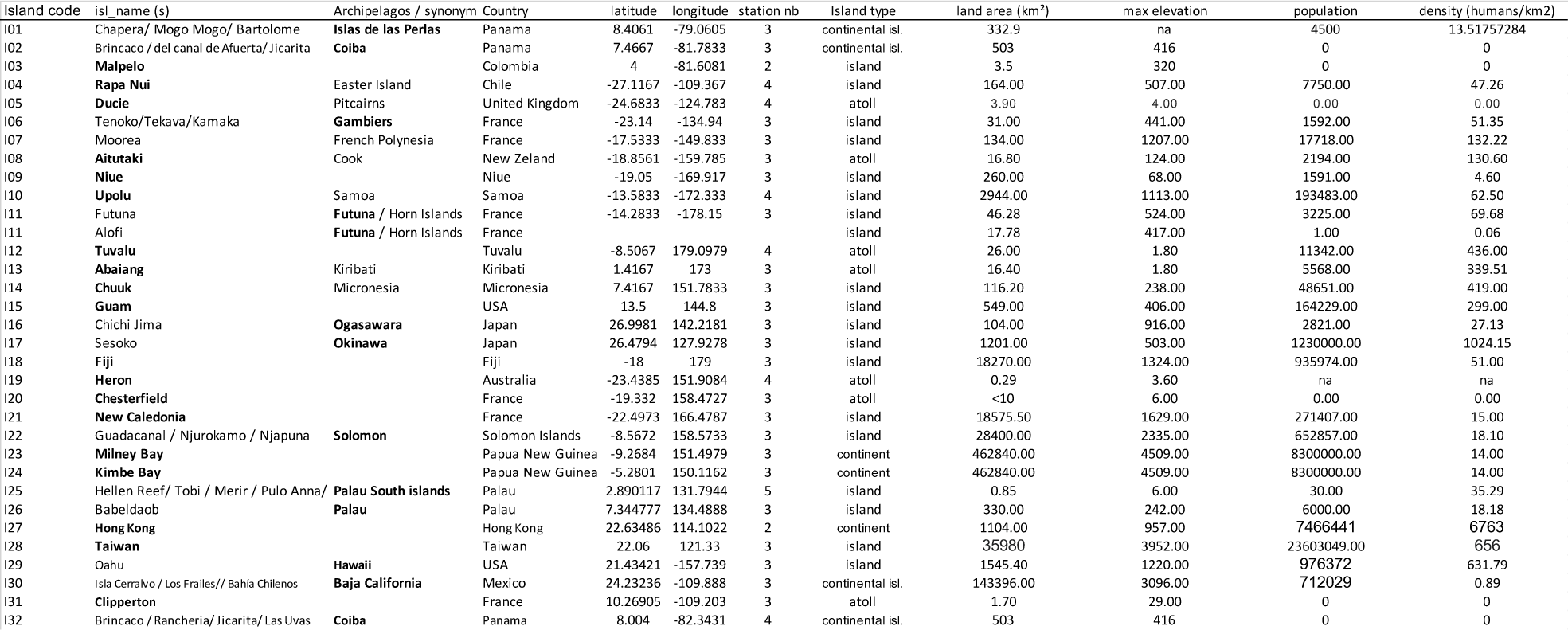
Summary of the different islands sampled during the Tara Pacific expedition with the associated island code (I01 to I32), their chosen reference name (in bold) corresponding either to the name of the island or of the archipelagos and some island characteristics recovered from various sources.

### 2 Sampling coral reef systems

The sampling event sequence and protocols were performed consistently over the whole expedition. Sampling was operated following the same procedure, approximate timing, and articulated around the same standardized “sampling events” (Figure 2) which allowed the same collection of samples with a standardized protocol (Table 2). On rare occasions, the timing and protocols were adapted for sailing conditions and to fit the schedule. Sampling events are characterized by their mode of sampling, which could be either directly from Tara’s dinghy [ZODIAC] or directly either using scuba-diving ([SCUBA]) or snorkeling ([SNORKEL]). In addition, the sampling device and strategy are included in the sample identifier.

**Figure 2:**
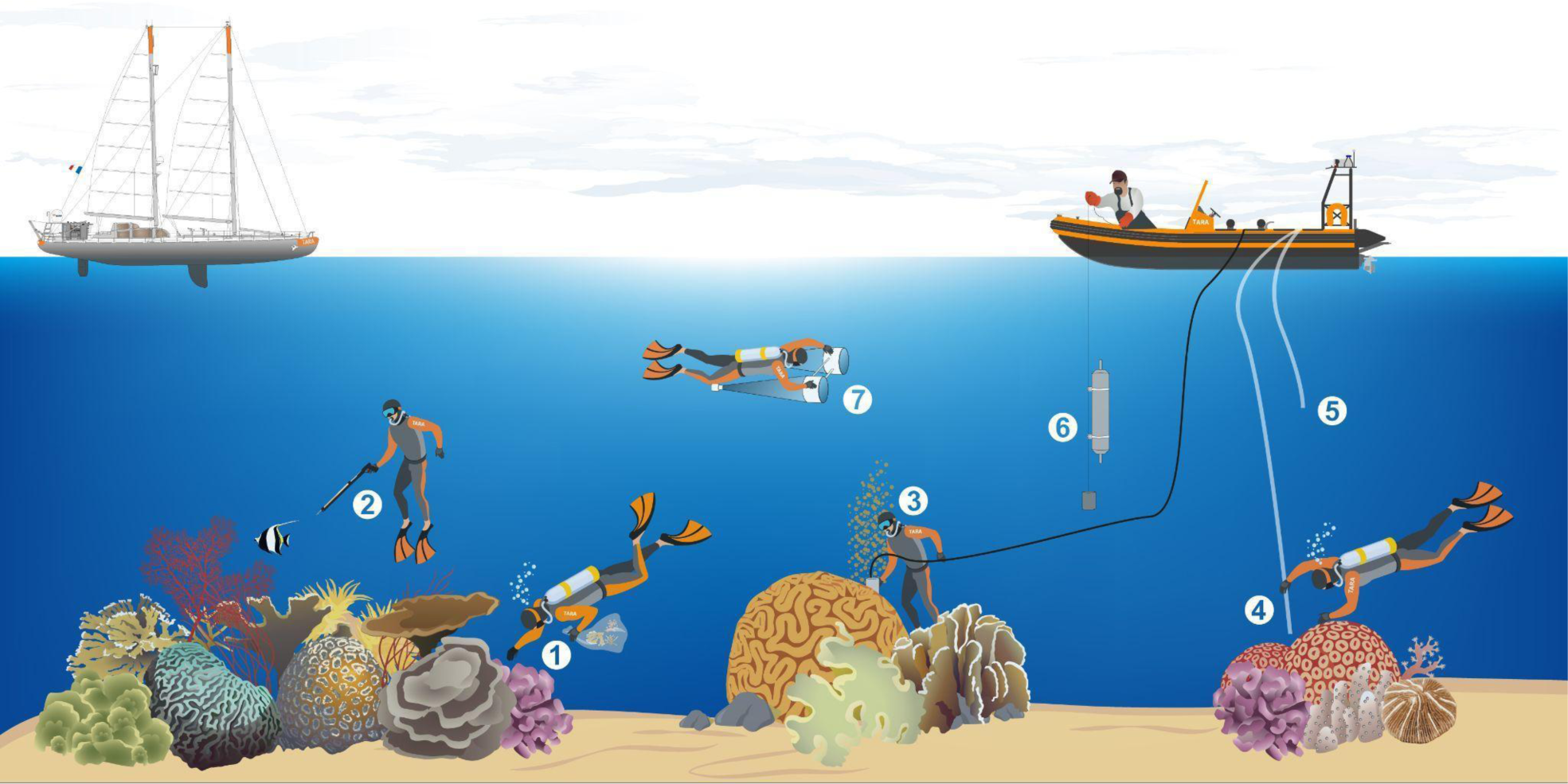
Schematic overview of the various sampling events conducted during the Tara Pacific expedition while sampling on coral systems. The different events are represented by the different numbers. (1) [SCUBA-3X10] and [SCUBA-SURVEY]; (2) [SNORKLE-SPEAR]; (3) [SCUBA- CORER]; (4) [SCUBA-PUMP]; (5) [ZODIAC-PUMP]; (6) [ZODIAC-NISKIN]; (7) [SCUBA- NET-20].

**Table 2.**
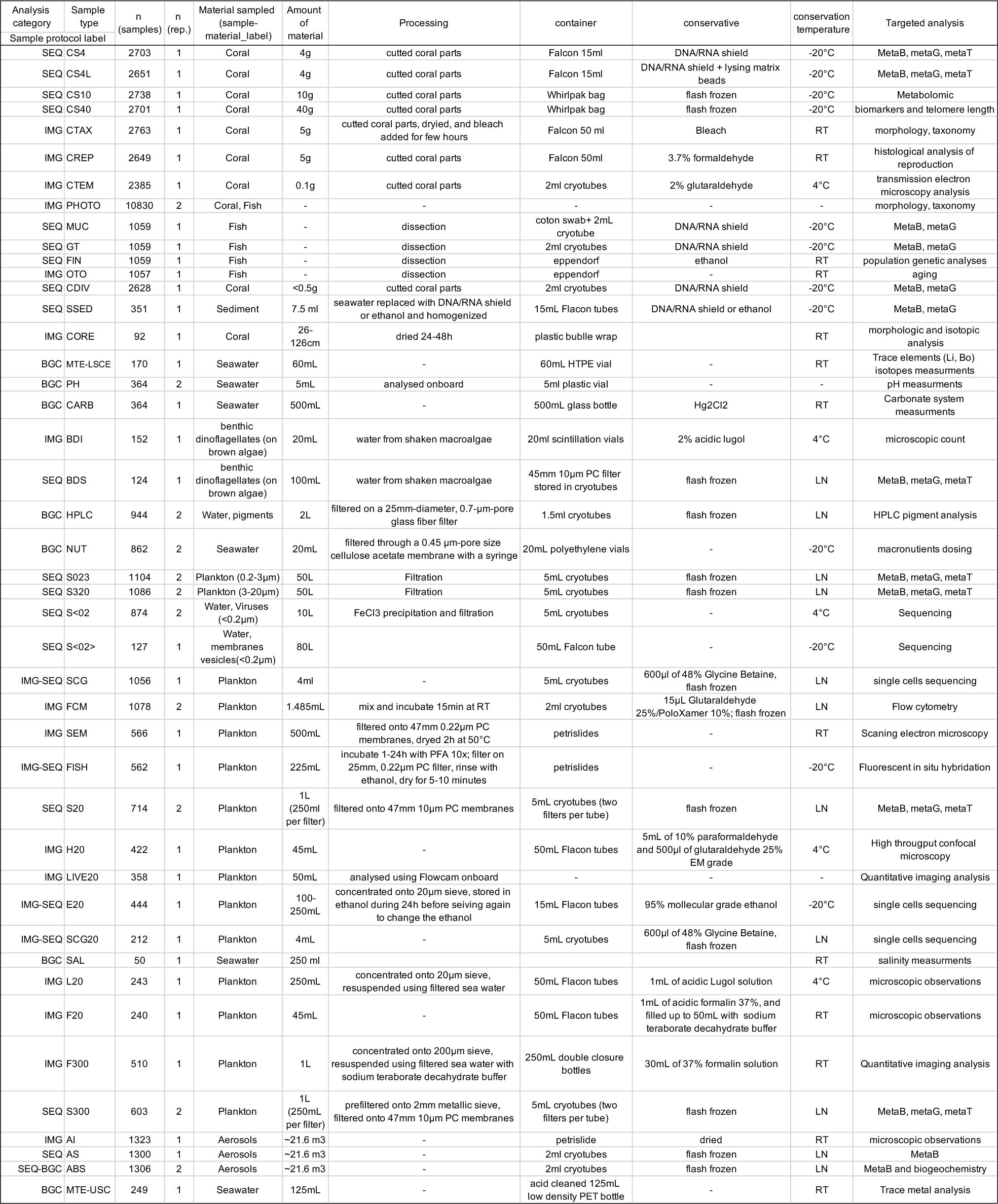
(online): Correspondence between samples types and their associated events and a summary of the protocol used and targeted analysis. RT: Room temperature, LN: Liquid Nitrogen, MetaB: metabarcoding, MetaG: metagenomic, MetaT: metatranscriptomic, PC: Polycarbonate, PET: Polyethylene (online version here:https://docs.google.com/spreadsheets/d/1ChLbq9GbUUvHZCzihGjnxZvmBOUFvGnv/edit?usp=sharing&ouid=105928735891310184253&rtpof=true&sd=true).

The first set of sampling events (usually in the morning) was mostly devoted to the sampling event [SCUBA-3X10] to sample coral colony fragments. In the meantime, another team pumped underwater, with the [SCUBA-PUMP] to collect coral surrounding water ([CSW]), while the third team snorkeled to capture a total of 10-15 fish using a speargun ([SNORKLE-SPEAR]). A small CTD probe (Castaway CTD) was also deployed from the dinghy down to the reef (generally ∼5 to 10m) to record temperature and conductivity profiles.

The second set of sampling events (usually in the afternoon) was devoted to a survey of coral diversity ([SCUBA-SURVEY]) concurrently with sampling surface water for biogeochemistry ([ZODIAC-NISKIN]), plankton in the size-fractions smaller than 20 µm ([ZODIAC-PUMP]), and plankton in the size-fractions between 20 to 2000 µm ([SCUBA-NET-20]). Finally, over a last dive a coral core was recovered over a large colony of *Porites* or *Diploastrea* ([SCUBA-CORER]).

#### 2.1 Sampling coral colonies [SCUBA-3X10]

During this typical sampling event, a total of 30 coral colonies [C001] to [C030], including 10 colonies for each of the 3 targeted species (*Pocillopora meandrina, Porites lobata*, and *Millepora platyphylla*) were sampled. Each colony was first photographed ([PHOTO]) using a 20 cm quadrat as a scale, their depth recorded and then sampled to collect about 70 grams of each coral by mechanical fragmentation using hammer and chisel. Fragments were placed in Ziploc bags labelled by colony ID and brought back to the boat.

#### 2.2 Sampling coral surrounding water [SCUBA-PUMP] and [ZODIAC-NISKIN]

Two *Pocillopora meandrina* coral colonies [C001] and [C010] were marked with small buoys, and [CSW] samples were collected as close as possible to the coral colony before the actual SCUBA-3X10 sampling to avoid contamination of the water samples with fragments or tissues released during the mechanical fragmentation of coral colony. Then, water was pumped using a manual membrane pump onboard Tara’s dinghy that was stationary above the coral colony. A scuba diver was holding a clean water tubing next to the colony while the operator onboard the dinghy was pumping the water up to the skiff. First, the water collected was used to rinse the pumping system, as well as a 20 µm metallic sieve and the 50 L carboys that will be used to transport the sample [C010]. Then, 50 L of water was filtered within and around the coral colony onto a 20 µm metallic sieve and directly stored in the dedicated clean 50 L carboy ([SCUBA-PUMP] for [C010]). When available, two replicates of sediment samples (i.e. sand [SSED]) were also taken using two 10 mL cryovials near the sampled colony. Finally, the coral colony [C010] itself was sampled following the [SCUBA-3X10] protocol.

Once the [C010] sampled, the dinghy was moved on top of colony [C001], where, before any other sampling was performed, carbonate chemistry and nutrients protocols (using a 5L Niskin bottle for carbonates [CARB] and nutrients [NUT]) as well as for [PH] protocols (using 5mL polypropylene vials and a 50mL Falcon tube) were performed. The [PH] was first sampled using two vials (5 mL polypropylene vials for samples), and a falcon 50mL tube (for later use to rinse the probe) were first lowered closed, opened next to the colony, rinsed with the [CSW], and closed tightly making sure no bubbles were trapped inside the vials. Next, the Niskin bottle was immersed open by the diver [ZODIAC-NISKIN], well rinsed along the descent and with the coral surrounding water near the targeted colony, and finally closed as close as possible from the colony [C001]. The tubing, the sieve, a 4L Nalgene (protected with reflective tape to isolate the sample from sunlight), and the 50L carboy dedicated for [C001] were rinsed with the [C001] [CSW]. The 5L Nalgene bottle was filled with [C001] [CSW] for high-performance liquid chromatography (HPLC). The 50L carboy was then filled ([SCUBA-PUMP] for [C001]) and the sediment samples [SSED] were collected following the same procedure as for [C010]. Finally, the coral colony [C001] itself was sampled following the same [SCUBA-3X10] protocol. For safety reasons, carbonate chemistry samples [CARB] could not be preserved with mercury (II) chloride on-board the dinghy due to its acute toxicity. Hence, the Niskin bottle was sampled on the last colony of the sampling sequence to minimize the time between sampling and chemical preservation on-board Tara.

#### 2.3 Sampling for fish [SNORKLE-SPEAR]

Fish sampling of two target species (*Acanthurus triostegus* and *Zanclus cornutus*) was operated by spear-fishing and snorkeling for a target number of about 10-15 fishes ([F001] to [Fxxx]) depending on the population present. The targets were speared and immediately stored in labeled individual Ziplock bags to avoid contamination between samples and kept inside a floating container to keep them at water temperature.

#### 2.4 Sampling sediments and macroalgae [SCUBA-…]

Sediments and macroalgae samples were sampled when encountered during the different dives. Sediment samples (i.e. sand [SSED]) were taken using two 10 mL cryovials near the sampled colony. Macroalgae, ideally brown macroalgae with thallus morphology type arbustive, ([MA01]- [MAxx]) were photographed ([PHOTO]) and sampled in individual Ziplock bags when encountered.

#### 2.5 Coral biodiversity sampling [SCUBA-SURVEY]

Biodiversity sampling transects were conducted in two depth-range environments to sample up to 80 coral colonies ([C041] to [C120]) randomly chosen with ideally up to 40 colonies living at a depth of 10-16 m, and up to 40 colonies living at a depth of 2-10 m, with an emphasis on sampling across a diverse range of coral hosts at different depth. Two pictures of each colony sampled were taken ([PHOTO]), and small pieces of 1-3 cm^2^ were sampled using a hammer and a chisel or a bone cutter.

#### 2.6 Sampling surface seawater [ZODIAC-NISKIN] and [ZODIAC-PUMP]

In addition to the seawater collected next to coral colonies explained above, surface ([SRF]) seawater was sampled at 2 m depth using the manual pump on-board of the dinghy ([ZODIAC- PUMP]). The [SRF] site was chosen to be as close as possible from the coral colonies sampled in the morning but with enough water depth that the plankton net sample could be taken at 2 m depth and at least 5 m above the seafloor. When the sampling site was shallower than 7 m, the site was chosen where these sampling conditions could be met within 100 m around the [CSW] sampling site. The water collected was treated similarly to the [SCUBA-PUMP] samples, with the difference that 100 L [SRF] water was collected into two 50 L carboys. The 4 L Nalgene bottles protected from sunlight were also filled with water at 2 m below the dinghy for HPLC filtrations on-board Tara.

#### 2.7 Sampling large size plankton [SCUBA-NET-20]

During this surface water pumping, plankton larger than 20µm were sampled at 2m below the sea surface using two small diameter bongo plankton nets with 20µm mesh size, attached to an underwater scooter ([SCUBA-NET-20]) and towed for about 15min at maximum speed (0.69 ± 0.04 m.s^-1^). The average maximum speed of the net tow was estimated in Taiwan (island 28 site 03) measuring the time it took the diver with full gears on and the nets attached, to travel between two buoys separated by a 9-meters line held tight and floating with the current, to avoid any impact of the current. The measurement was repeated three times facing the current, three times in the same direction as the current, and five times with the current sideways. Each net was equipped with flowmeters, but the speed of the underwater scooter was insufficient to trigger their rotation, therefore the time of sampling was precisely timed to estimate theoretically the volume filtered using the following equations:

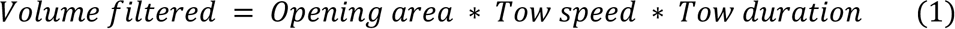

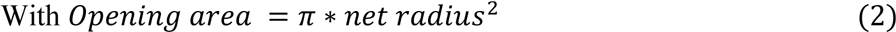

The volume estimated from the flowmeter reading was about 60 times smaller than the volume calculated theoretically, thus, only the theoretical volume will be used in concentration calculations.

After 15 minutes of towing, the divers surfaced the two nets and the two cod-ends were sieved through a 2000 µm metallic sieve, into a 2 L Nalgene (r) bottle. The bottle was topped-up with 0.2 µm filtered seawater from the same sampling site and kept at ocean temperature in a bucket during transportation to Tara. Finally, [PH], [NUT] and [CARB] samples were taken at 2 m depth just before leaving the sampling site following the same protocol than for [CSW] sampling and using the same cleaned 5 L Niskin bottle for [CARB] and [NUT], and two 5 mL polypropylene vials as well as a Falcon 50 mL tube for [PH].

#### 2.8 Sampling coral cores [SCUBA-CORER]

During the last dive, coral cores were sampled ([SCUBA-CORER]) on *Porites* colonies previously identified and photographed ([PHOTO]). To prevent contamination with coral fragments and tissues released during coring, two [CARB] samples of seawater were taken (one at the surface and one close to the coral colony) before coring and using two 500 mL glass stoppered bottles. Grease was applied to the glass stopper before the dive to allow opening under pressure next to the coral colony. The diver lowered the bottles closed, opened one at 2 m below the surface, and one next to the coral colony. Another seawater sample was taken with a 60 mL HDPE plastic bottle at 2m depth for subsequent analysis of trace isotopes in relation to the core analysis. Once all seawater was sampled, a 250 mm diameter, 600 round per minute corer from Melun Hydraulique was used to coral cores ([CORE]). Forty coral skeletal cores (40 – 150 cm long) were collected from colonies living between 3 m (Moorea Island-I07) and 20 m (Futuna Islands-I11) depth. From island I19 (Great Barrier Reef) the same protocols were also carried out on large *Diploastrea heliopora* colonies when encountered.

#### 2.9 Samples processing

##### 2.9.1 Benthic samples

Once back onboard Tara, the material collected during each sampling event was immediately processed into various samples. Samples were labeled with their target analysis (e.g. sequencing ([SEQ]), imaging, microscopical or morphological inspection ([IMG]) or biogeochemical measurements ([BGC])).

Coral samples obtained from [SCUBA-3X10] events were immediately sorted and separated using bone cutters, in several sub-samples usually labeled with the amount of material used or with the targeted analysis (Table 2). [CS4] and [CS4L] samples containing ∼4 g of coral material, were stored at -20°C in 15 ml Falcon tubes and 6 ml of DNA/RNA shield (Zymo Research, Irvine, CA, USA) for subsequent metabarcoding, metagenomic and metatranscriptomic analyses. [CS4L] only differs from [CS4] by the addition of lysing matrix beads. [CS10] and [CS40] samples, that contain respectively 10 g and 40 g of coral material, were stored in Whirlpak® sample bags, immediately flash frozen in liquid nitrogen, and kept at -20°C. These samples are intended for subsequent metabolomic analysis for [CS10], physiologic/stress biomarkers (symbiont and animal biomasses, antioxidant capacity and protein damages) and telomeric DNA length for [CS40]. Morphological taxonomic identification [CTAX] samples were performed by drying 5 g of material in 50 ml Falcon tubes, and removing organic material with the addition of 3-4% bleach solution during approximately 2 days. After discarding the bleach solution, clean skeletons were preserved dry at room temperature. For histological measurements of reproduction status [CREP], 5 g of each coral colony was preserved in a 50 ml Falcon tube filled with a 3.5% formaldehyde solution and stored at room temperature. Finally, for transmission electron microscopy examination of coral intracellular details including viruses [CTEM], 0.1 g of coral tissue was preserved with 250 µL 2% glutaraldehyde and conserved at 4 °C in a fridge.

Macroalgae samples ([MA]), and the seawater collected with them, were firmly shaken to resuspend attached epiphytic organisms. 20 mL of water was transferred into glass vials and fixed with 2% acidic Lugol and stored at 4 °C for future benthic dinoflagellates identification and counts using microscopy ([BDI]), while 100 mL of each replicate were filtered onto a 10 µm pore size polycarbonate filter which was flash frozen and preserved in liquid nitrogen for future metabarcoding analysis ([BDS]).

[SSED] samples were immediately flash frozen when brought back on-board Tara.

About 30 to 40 mL of the seawater that was sampled with the coral fragments of [C001] and [C010] and transported in the coral individual Ziplock bags were transferred immediately after the dive into 50 mL falcon tube and stored at water temperature in non-direct ambient light to recover cultures of plankton species closely associated with coral colonies ([IMG-LIVE]).

When fish were recovered onboard, a [PHOTO] was taken, their sex and length were determined before taking a sample of skin mucus ([MUC]) by collecting 1 cm^2^ of skin. The fish were then dissected to recover about 3 cm long of the final section of the digestive tract ([GT]) that was preserved in 2 mL cryotubes with 1 ml of DNA/RNA shield and then stored at -20 °C for metagenomic and metabarcoding analyses. One fin sample ([FIN]) was dissected, and preserved into an Eppendorf tube filled with 95° ethanol for population genetic analyses. Last, the otolith ([OTO]) was also dissected and stored dry into an Eppendorf tube at room temperature for later aging of each fish.

Coral samples obtained from [SCUBA-SURVEY] were collected for symbionts and coral diversity analysis ([CDIV]) using different marker genes (metabarcoding, 18S, 16S and ITS2).

About 0.5 g of material was preserved with DNA/RNA shield and stored into 2 mL cryotubes at -20°C.

Finally, samples collected during [SCUBA-CORER] events were also processed and stored onboard Tara. The [CORE] were rinsed with freshwater, air dried for 24-48h before being wrapped into a plastic bubble wrap for sclerochronological and geochemical analysis, to recover historical water biogeochemical properties. The [PH], [CARB] and [MTE-LSCE] samples associated with the coral core [CORE] were processed following the same protocol than the water samples collected with the [SCUBA-PUMP] and [ZODIAC-PUMP] (explained in section 2.2.2), with the exception that the [CARB] and [MTE-LSCE] samples were already stored in their final container during sampling on the dinghy.

##### 2.9.2 Water samples for biogeochemistry

The [PH] was measured from the two replicates 5 mL polypropylene vials onboard Tara using an Agilent Technologies Cary 60 UV-Vis Spectrophotometer equipped with an optical fiber. The detailed protocol was previously described^13^, but briefly, the 5 mL vials and the 50 mL falcon tube were kept closed and acclimated to 25°C for 2–3 h. Absorbance at specific wavelengths was then read before and after the addition of 40 µL meta-Cresol Purple dye to each 5 mL vial. The probe was rinsed between each measurement using the 50 mL falcon tube containing the same seawater as the 5 mL vials samples. TRIS buffer solutions^17^ were measured regularly along the cruise to validate the method and correct for potential drifts of pH of the dye solution.

The Niskin bottles of the morning ([CSW] for [C001] colony) and afternoon ([SRF]), carefully kept closed since sampling on the dinghy, were each used to rinse and fill one 500 mL glass stoppered bottle on Tara. Some grease was applied to the glass stopper, and bottles were filled with water samples leaving 2 mm of air below the bottom of the bottleneck. Note that the [CARB] samples associated with the [CORE] samples were already stored in their final container and grease was already applied to the glass stopper before the dive. The water level of these samples was simply adjusted to 2 mm below the bottleneck. All [CARB] samples were immediately poisoned with 200 µL of saturated mercury (II) chloride solution (HgCl2) and stored at room temperature.

The Niskin water was then used to rinse and fill up trace element samples in 60mL HDPE plastic bottles [MTE-LSCE]. These samples were stored at room temperature and used to confirm the absence of local influence on Li and B isotopic signals. Similar to [CARB] associated with [CORE] samples, the [MTE-LSCE] samples associated with [CORE] samples were already stored in their final containers, therefore, were just stored at room temperature.

The water remaining from the Niskin bottle, sampled in the morning ([CSW] for [C001] colony) and the afternoon ([SRF]), was used to prepare macronutrient samples ([NUT]). A 50 mL syringe was rinsed with the sampled seawater three times. A filter 0.45 µm-pore size cellulose acetate membrane was then connected to the syringe and ∼20 mL of sample water was run through it to rinse the filter. Once the syringe, filter and vials were properly rinsed twice, two 20 mL polyethylene vials were filled running the sampled water through 0.45 µm-pore uptidisc syringe filter. Nutrient samples were stored vertically at -20°C.

Two replicates of two liters of seawater sampled in the 4L Nalgene bottle from the [SCUBA- PUMP] and [Zodiac-PUMP] events, were filtered onto 25mm-diameter, 0.7-µm-pore glass fiber filters (Whatman GF/F) and immediately stored in liquid nitrogen for later High-Performance Liquid Chromatography ([HPLC]) analysis to obtain pigments concentration.

##### 2.9.3 Water samples for genomics and imagery

The water collected during the [SCUBA-PUMP] and [Zodiac-PUMP] events was treated similarly, with the only difference that while [Zodiac-PUMP] samples were treated in duplicates, the two 50 L samples collected during [SCUBA-PUMP] correspond to [C001] and [C010] colonies. This applies only for sequencing samples ([SEQ-S]), while all other samples were taken in duplicates. Additionally, all genomic samples were processed to be as comparable as possible with previous existing samples from Tara Oceans^11, 14^.

As soon as back on-board Tara, the water collected was used to rinse and fill one (for each [CSW]) or two (for [SRF]) 50 L carboy but also to fill two 2L Nalgene(r) bottles. The content of the 50 L carboys was immediately size-fractionated by sequential filtration onto 3-µm-pore-size polycarbonate membrane filters and 0.22-µm-pore-size polyethersulfone Express Plus membrane filters. Both were placed on top of a woven mesh spacer Dacron 124 mm (Millipore) and stainless-steel filter holder “tripods” (Millipore). Water was directly pumped from the 50 L with a peristaltic pump (Masterflex), and separated into samples that contain particles from 3-20µm ([S320]) and 0.2-3µm ([S023]) for latter sequencing. To ensure high-quality RNA, the filtering of the first replicate ([C001] for [CSW] samples and any of the two 50 L carboys for [SRF]) were stopped after

15 min of filtration while the second was continued for the full volume (or a maximum of 60 min) to maximize DNA yield. Filters were folded into 5 mL cryovials and preserved in liquid nitrogen immediately after filtration. During this filtration 10 L of 0.2 µm filtered water ([S<02]) was collected from each replicate, 1 mL of FeCl3 solution was added to flocculate viruses^18^ for 1 hour. This solution was then again filtered onto a 1-µm-pore-size polycarbonate membrane filter using the same filtration system as for [S320] [S023]. Filters were then stored in 5 mL cryotubes and stored at 4° C for later sequencing of viruses. The 80L remaining of 0.22 µm prefiltered water was used to filter membranes vesicles ([S<02>]) using an ultrafiltration Pellicon2 TFF system by keeping the pressure below 10 psi until the concentrate was reduced to a final volume of 200-300 mL. This sample was further concentrated using a Vivaflow200 TFF system at a recirculation rate of 50-100 mL/min and less than one bar of pressure until obtaining a final sample of 20mL. Flushing back the system usually brings this volume to up to 40mL which was stored in a 50 mL Falcon tube at -20°C.

Two 4mL samples were taken from the 2 L Nalgene bottles, and stored into 5 ml cryotubes fixed with 600μl of 48% Glycine Betaine and directly flash-frozen for later single cells genomic analysis ([SCG]). For flow cytometry cell counting ([FCM]), two replicates of 1.485 mL of sampled water were placed into 2 mL cryotubes pre-aliquoted with 15 μL of fixative composed of Glutaraldehyde (25%) and PoloXamer (10%). Tubes were then mixed gently by inversion, incubated 15min at room temperature in the dark before being flash-frozen, and kept in liquid nitrogen. For scanning electron microscopy ([SEM]), 500mL of water was filtered onto a 47mm 0.22µm pore size polycarbonate filter, placed in a petri slide, dried for two hours at 50°C and conserved at room temperature. Fluorescence In Situ Hybridization ([FISH]) samples were prepared by adding 225 mL of seawater into a 250 mL plastic vial containing 25 ml of 10xPFA. The samples were incubated at 4°C before filtration onto two 25 mm 0.22 µm pore size polycarbonate filters, rinsed with ethanol, placed in petri slides, dried for 5-10minutes before being stored at -20°C.

Samples collected during the [SCUBA-NET-20] were processed to obtain different samples for sequencing and imaging needs. One litre of the sample collected was filtered onto four 47mm 10µm pore size polycarbonate membranes (250mL each). Filters were then placed into 5mL cryotubes, flash-frozen, and stored in liquid nitrogen for later sequencing ([S20]). 45 mL was subsampled into a 50ml Falcon tube, fixed with 5mL of 10% paraformaldehyde and 500μl of glutaraldehyde 25% EM grade, and stored at 4°C for future high-throughput confocal microscopy ([H20]; e.g.^19^). 4mL was stored in 5mL cryotubes, fixed with 600µl of 48% glycine betaine, immediately flash frozen and kept in liquid nitrogen for single cell genomics ([SCG20]). Another sample for single cell sequencing stored in ethanol ([E20]) was done by filtering 100 to 250 mL of the sample onto a 20µm sieve and re-suspended in EM grade ethanol for 24h at 4°C. After incubation, the sample was sieved a second time to remove any trace of seawater, re-suspended with EM grade ethanol into 15 mL falcon tube, and stored at -20°C. Finally, a 50mL sample was directly imaged live onboard ([LIVE20]) using a Flowcam^20^ Benchtop B2 series equipped with a 4x lens and processed using the auto-image mode.

### 3 Oceanic sampling

To obtain both a large scale and local (around coral reef island) environmental characterization, a comprehensive set of physical, chemical and biological properties of the sea surface ecosystem was recorded while cruising. This sampling scheme was framed to be compatible with the previous Tara Ocean expedition measurements^11, 14^, but also to provide a continuity with water samples conducted directly on the coral reef. Furthermore, while the biology and ecology of surface ecosystems remain largely unknown, they are an essential component of air-land-sea exchanges and are subjected to numerous hydrological, atmospheric, physical and radiative constraints^21^ and is therefore at the frontline of climate change and pollution.

The main goals and general overview of this sampling are already described^13, 22^ and will be briefly presented here in the context of the different sampling events and samples that were generated. Measurements and samples could be separated into two types: i. local samples originating from a local sampling event, and ii. autonomous high frequency continuous measurements of atmospheric and surface seawater properties (e.g., per minute averages of higher frequency measurements). In the case of the discrete water sampling, the different sampling events were either attributed to a station (noted [OA001] to [OA249]) if they were conducted in a reasonably short time lapse (> 75 km away, or > 0.25 days away from a group of OA Events), or noted [OA000] otherwise. Similarly, every OA station located within 200 nautical miles (370 km) of an island were annotated with that Island label, i.e. the sampling-design-label of the corresponding OA Events and OA Samples is [OA###-I##-C000]. The continuous sampling was conducted as follows: a. surface seawater measurements were performed by pumping water continuously through the boat hull ([INLINE-PUMP]) at ∼1.5 m depth, b. light and atmosphere properties were measured 5 m above the sea level ([PAR + BATOS]), and c. aerosols were sampled by pumping air on top of the mast ([MAST-PUMP]) at ∼27 m (15 m during the first trans-Atlantic transect prior to May 2016).

#### 3.1 Sampling events

Sampling was organized following several successive events, generally at daily frequency, in the morning. Water collection while cruising was carried out by a custom-made underway pumping system nicknamed the [DOLPHIN] connected by a 4 cm diameter reinforced tubing to a large volume industrial peristaltic pump (max flow rate = 3 m^3^ h^-1^) on the deck. The system was equipped with a metallic pre-filter of 2 mm mesh size, two debubblers, and a flowmeter to record the volume of water sampled. Unfiltered water was collected first for a series of protocols, water was prefiltered using a 20 µm sieve to rinse and fill two 50 L. Both unfiltered seawater use and 20µm filtered seawater were labelled as [CARBOY]. To collect larger plankton, water was pumped from the DOLPHIN into a 20 µm net fixed on the wetlab’s wall ([DECKNET-20]) for 1 to 2 hours depending on biomass concentration simultaneously to a net tow using a “high speed net” ([HSN-NET-300]). The HSN was equipped with 300 µm mesh sized net and designed to be efficient up to 9 knots. It was towed from 60 to 90 minutes depending on the plankton density. Near islands and in the Great Pacific Garbage Patch, a Manta net ([MANTA-NET-300]) with a 0.16 x 0.6 m mouth opening with a 4 m long net with 300 µm mesh size was used concurrently at a maximum speed of 3 knots. Finally, trace metal samples ([MTE-USC]) were collected from the bow using a metal-free carbon fibber pole [HANDHELD-BOW-POLE] on which a plastic fixation have been added to insert a 125mL low density polyethylene bottle (LDPE) which was previously pre-washed on land and stored individually in separate ziploc bags. To avoid contamination from the boat, samples were hand held collected, wearing polyethylene gloves, while cruising upwind on the bow of the boat (i.e., before the boat got in contact with the collected water; Figure 3).

**Figure 3:**
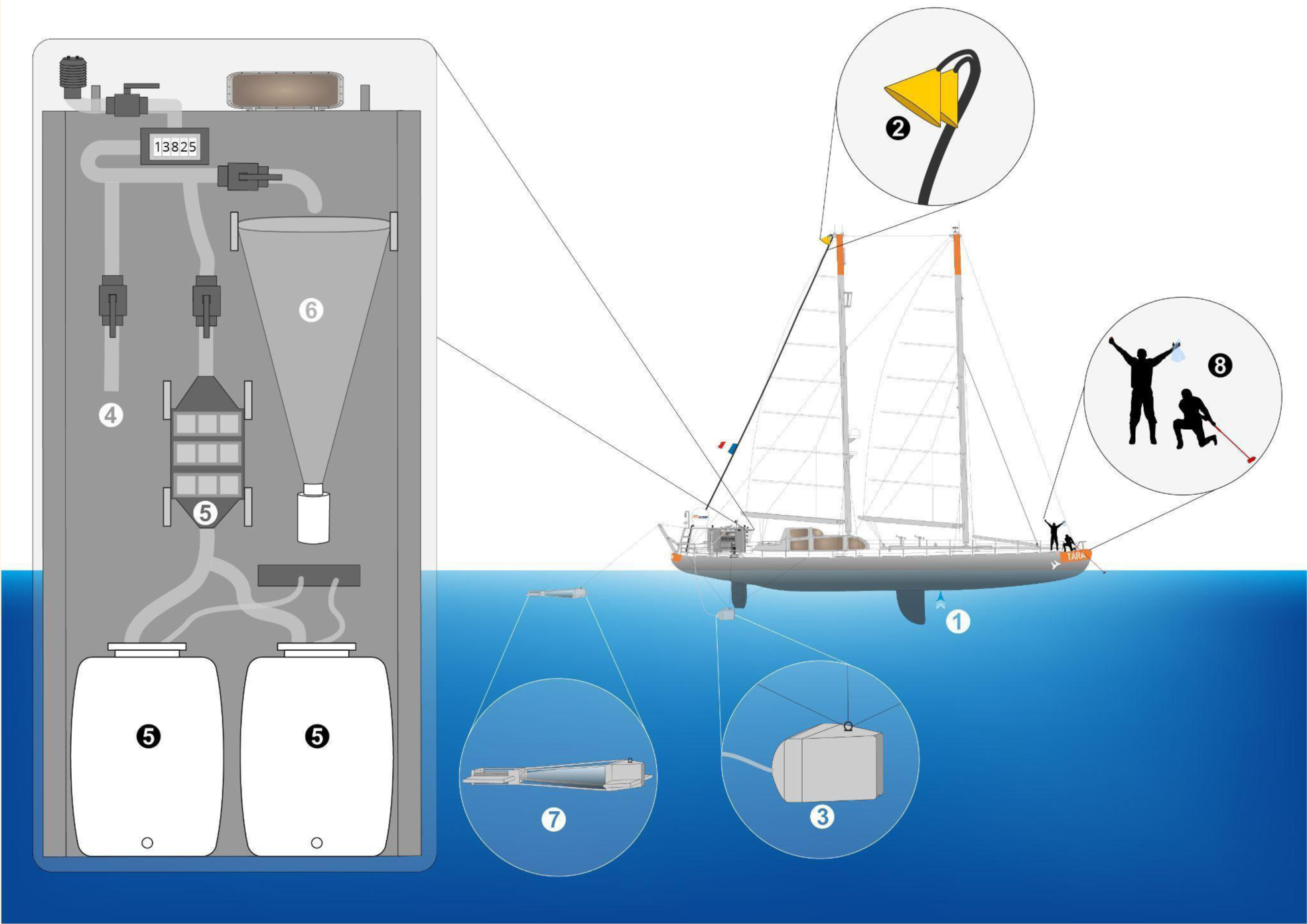
Schematic overview of the various sampling events conducted during the Tara Pacific expedition while sampling on oceanic stations. The different events are represented by the different numbers. (1) [INLINE-PUMP]; (2) [MAST-PUMP]; (3) [DOLPHIN] pumped water that is either used (4) [RAW], filtered at 20µm to fill two 50L (5) [CARBOY], or filtered though (6) [DECKNET-20]; (7) “high speed” [HSN-NET-300] or [MANTA-NET-300] plankton nets; (8) [HANDHELD-BOW-POLE].

#### 3.2 Samples processing

Water, plankton and aerosols samples collected in the vicinity of islands and from the open sea were processed as much as possible following similar protocols than on islands. Samples collected both on islands and in open sea are marked with asterisks* here, and only the few differences in protocols will be noted.

##### 3.2.1 From Dolphin, unfiltered water

Unfiltered seawater collected from the [DOLPHIN] was used to process several samples for biogeochemical purposes ([BGC]). For every station, samples were collected for nutrients [NUT]*, [PH]* measurements and pigments analysis by [HPLC]*. Salinity [SAL], carbonates ([CARB]*) and trace elements [MTE-LSCE]* were sampled on a weekly basis. [SAL] samples were done by sampling 250 mL of seawater in a 250 mL hermetically sealed glass bottle.

##### 3.2.2 From Dolphin, pre-filtered water

The two 50 L carboys of 20 µm prefiltered seawater were used to produce size fractionated samples for genomic analyses ([S320]* [S023]* [S<02]*). The same pre-filtered seawater was sampled for flow cytometry cell counting ([FCM]*) and single cell genomic ([SCG]*).

##### 3.2.3 From Dolphin-Decknet

Once the [DECKNET-20] time limit reached (between 1 and 2 hours), the flow was stopped and the net was carefully rinsed with 0.2 µm filtered seawater. The plankton sample was then transferred to a 2 L Nalgene bottle and completed to 2 L with 0.2 µm filtered seawater. The sample was homogenized by repeated smooth bottle flips and split into four 250mL subsamples for [S20]*, one 250 mL sample for [E20]*, one 250 mL sample for [LIVE20]*, and one 45 mL sample for [H20]*. In addition to these already described protocols, one 250 mL sample was also taken for [L20], for which the seawater was drained using a 20µm sieve and the plankton was transferred in a 50 mL Falcon tube and fixed with 1 mL of acidic lugol solution for latter microscopic observations. Finally, a 45 mL sample was taken for [F20], transferred in a 50 mL Falcon tube and fixed with 1 mL of 37% formalin solution and completed to 50 mL with sodium tetraborate decahydrate buffer solution for latter microscopic observations.

##### 3.2.4 From HSN/Manta nets

Once recovered, samples collected both by the HSN net and the Manta net followed the same procedure. The net was carefully rinsed from the exterior to drain organisms into the collector. Its content was transferred using 0.2µm filtered sea water in a 2L Nalgene Bottle and completed to 2L. The sample was then homogenized and split in two 1L samples. The first half was prefiltered onto a 2mm metallic sieve and filtered onto four 47mm 10µm pore size polycarbonate membranes (250mL each), Filters were then placed into 5mL cryotubes, flash frozen and conserved in liquid nitrogen for latter sequencing ([S300]). The second fraction was concentrated onto a 200µm sieve and resuspended in a 250mL double closure bottle using filtered seawater saturated with sodium tetraborate decahydrate, fixed with 30mL of 37% formalin solution and stored at room temperature for latter taxonomic and morphological analysis using imaging methods ([F300]).

##### 3.2.5 From Mast-pump

Aerosols pumped through one of the ([MAST-PUMP]) inlets were channelled through a conductive tubing of 1.9 cm inner diameter to four parallel 47mm filter holders installed in the rear hold using a vacuum pump (Diaphragm pump ME16 NT, VACUUBRAND BmbH & Co KG, Wertheim, Germany) at a minimum flow rate of 30 lpm (20lpm prior to May 2016). Three filter holders were equipped with 0.45µm pore size PVDF filters for latter aerosol sequencing ([AS]) and biogeochemical analysis together with sequencing ([ABS]), while the fourth one was a 0.8µm pore size polycarbonate filter for later aerosol imaging ([AI]) analysis using scanning electron microscope. Twice a day (12h pumping periods), at approximate dusk and dawn, those filters were changed, [AS] and [ABS] filters were placed into 2mL cryotubes (2 filters for each [ABS] sample) and immediately flash frozen while [AI] filters were packaged in sterile PetriSlide preloaded with absorbent pads and stored dry at room temperature.

#### 3.3 Continuous measurements

As previously described (see^13, 22^), a comprehensive set of sensors were combined to continuously measure several properties of the water but also atmospheric aerosols and meteorological conditions. All sensors were interfaced to be synchronized with the ship’s GPS and synchronized in time (UTC time). Surface seawater was pumped continuously through a hull inlet located 1.5 m under the waterline using a membrane pump (10 LPM; Shurflo), circulated through a vortex debubbler, a flow meter, and distributed to a number of flow-through instruments. A thermosalinograph [TSG] (SeaBird Electronics SBE45/SBE38), measured temperature, conductivity, and thus salinity. Salinity measurements where intercalibrated against unfiltered seawater samples [SAL] taken every week from the surface ocean, and corrected for any observed bias. Moreover, temperature and salinity measurements were validated against Argo floats data collocated with Tara. A CDOM fluorometer [WSCD] (WETLabs), measured the fluorescence of coloured dissolved organic matter [fdom]. An [ACS] spectrophotometer (WETLabs) measured hyperspectral (4 nm resolution) attenuation and absorption in the visible and near infrared except between Panama and Tahiti where an AC-9 multispectral spectrophotometer (WETLabs) was used instead. A filter-switch system was installed upstream of the [ACS] to direct the flow through a 0.2

µm filter for 10 minutes every hour before being circulated through the [ACS] allowing the calculation of particulate attenuation [ap] and absorption [cp], by removing the signal due to dissolved matter, drift, and biofouling^23^. From November 13, 2016 to May 6, 2017, a backscattering sensor [BB3] (WETLabs ECO-BB3) in a flowthrough chamber (BB-box) was added to the underway system, upstream of the switch system, to measure the volume scattering function [VSF] at 124° and 3 wavelengths (470, 532, 650 nm) and estimate the backscattering coefficient [bbp]. From May 7^th^ 2017 to the end of the expedition, the BB-box and the [BB3] were moved downstream of the filter-switch system to run 0.2 µm filtered seawater for 10 minutes every hour in order to remove the biofouling signal and improve [bbp] estimations. Chlorophyll a content [chl] was estimated from [ap]^24^ and [cp] (when [cp] is hyperspectral^25^), as well as other pigments (when [ap] is hyperspectral^26^). The [chl] estimated from [ap] was then calibrated against the [HPLC] [chl]^24^. The particulate organic carbon concentration [poc] was estimated both using an empirical relation^27^ between measured [poc] and measured [cp], or applying an empirical relation between measured [poc] and [bbp]^28^. Phytoplankton organic carbon [cphyto] was estimated by an empirical relationship with [bbp]^29^. An indicator for size distribution of particles between 0.2 and ∼20 µm [gamma] was calculated from [cp]^30^. A brief description of the methods to analyse, calibrate, correct, and estimate bio-optical proxies are detailed in the section Technical Validation and more extensively explained in each processing report attached with the dataset.

An Equilibrator Inlet Mass Spectrometer [EIMS] (Pfeiffer Vacuum Quadrupole 1–100 amu) measured the Oxygen to Argon ratio in percent [o2ar], coupled with an optode (Aanderaa optode 4835) measuring oxygen concentration in the seawater [O2]. Concurrently with samples collected through the [MAST-PUMP], two instruments were installed aboard Tara to measure the size distribution and abundance of atmospheric aerosol particles: a scanning mobility particle sizer ([SMPS], SMPS-C GRIMM Aerosol Technik Ainring GmbH & Co. KG, Ainring, Germany) measuring particles in the size range 0.025 – 0.70 µm, and an optical particle counter ([EDM]; EDM180 GRIMM Aerosol Technik Ainring GmbH & Co. KG, Ainring, Germany) measuring all particles in the size range 0.25 – 32 µm. The SMPS was set to perform a full scan of particle distribution every 5 min and the EDM produced a particle size distribution every 60 s. Data provided from [EDM] includes both the total particle concentration (nb cm^-3^) in the size range 0.25 – 32 µm every 60 seconds, and through a second dataset averaged every 30 minutes, both the particle concentration (nb cm-3) together with its normalized size distribution (dN/dlogDp (nb cm^-3^), i.e., the concentration divided by the log of the size width of the bin),while data from [SMPS] are averaged at the hour scale and provided both at the scale of particle concentration (nb cm^-3^) together with its normalized size distribution (dN/dlogDp (nb cm^-3^)) .

Together with navigation data such as speed over ground [sog] and course over ground [cog] meteorological station (BATOS-II, Météo France) measured air temperature, relative humidity, and atmospheric pressure at 7 m above sea level. True and apparent wind speed and direction was measured at about 27 m above sea level. In October 2016 a Photosynthetically Active Radiation [par] sensor (Biospherical Instruments Inc. QCR-2150) was mounted at the stern of the boat (∼5 m altitude).

## Data Records

The full collection of datasets has been deposited either at Pangaea or at Zenodo depending on their nature, but also on the likelihood to be updated.

### Provenance metadata

Tara Pacific datasets are articulated around a consistent set of provenance metadata that provide temporal (UTC date and time) and spatial (latitude, longitude, depth or altitude) references as well as annotations about environmental features and place names, using controlled vocabulary from the environmental ontology (https://ebi.ac.uk/ols/ontologies/envo) and the marine regions gazetteers (https://marineregions.org/). These metadata are available at three granular levels: sampling stations and sites, sampling events, and samples collected at a specific depth.

A [sampling-design-label] is provided to facilitate the identification and integration of data that originate from the same open ocean station (OA###), island (I##), site (S##) or coral colony (C###), and hence share provenance and environmental context. For example, data originating from coral colony number twelve on the second site of the fourth island visited by Tara will bear the sampling design label OA000-I04-S02-C012. Similarly, data collected at station number 99 in the middle of the Pacific Ocean will bear the sampling design label OA099-I00-S00-C000, and data collected at open ocean station number 41 within 200 nautical miles of island number four will bear the sampling design label OA041-I04-S00-C000.

Each sample is also characterized by its sampling event which have several properties such as its date and time (UTC) of sampling ([sampling-event_date_time-utc]), the type of event from which the sample originates ([sampling-event_device_label]), the material sampled ([sample- material_label]; see Table 2), the protocol used ([sampling-protocol_label]; see Table 2) and finally the barcode attributed to the final sample obtained and replicated on the logsheets ([sample- storage_container-label]). Finally, each sample, in addition to its original barcode was characterized by an event label and a sample label composed of that previous information such as:

Sample label: TARA_SAMPLE_[sampling-event_date_time-utc]_[sampling-design label]_[sampling-environment_feature_label]_[sample-material_label]_[sampling- protocol_label]_[sample-storage_container-label]

Event label: TARA_EVENT_[sampling-event_date_time-utc]_[sampling-design label]_[sampling-day-night_label]_[sampling-environment_feature_label]_[sample- material_label]_[sampling-protocol_label]_[sample-storage_container-label]

The provenance context of all samples collected during the Tara Pacific Expedition is available as a single UTF-8 encoded tab-separated-values file, in open access at Zenodo and replicated in part at BioSamples (XYZ). In addition to georeferences and place names, the provenance metadata includes sample unique identifiers, taxonomic annotation from NCBI, and links to sampling logsheets and campaign summary reports.

Additionally, the full repository containing the campaign summary reports, sampling authorisations, logsheets and the full record of coral images could be consulted on Pangaea (https://store.pangaea.de/Projects/TARA-PACIFIC/). The full list of sampling events is consultable on the following repository: https://doi.pangaea.de/10.1594/PANGAEA.944511.

## Environmental context for data analysis

Rich collection of environmental parameters collected from either samples, on-board measurements, satellite imagery, operational models or even calculated from astronomical atlas were compiled and made available for further analysis. These environmental measurements were provided in a multi-layered way in open access to either Pangaea or Zenodo (Table 3), depending on the potentiality to require updates, with (1) raw measurements at the measure level for both physical samples or for on-board continuous measurements, accompanied with their quality check flags (2) a combined version regrouping all measurement at the sampling event level and adding satellite imaging and results obtained from operational models. (3) This latter was propagated, together with all measurements done on samples, to provide an environmental context to every collected samples belonging to the same station, but by also providing indices of the spatial ([dxy]), temporal ([dt]) and vertical ([dz]) discrepancies between the various measures and the designed sample and their variability (as assessed by mean, standard deviation, number of measures and 5, 25, 50, 75, 95 percentiles when possible); (4) a simplified version at the site level where all synonym measurements were cross-compared and chosen by level of quality. (5) At the scale of the site level, a series of Lagrangian and Eulerian diagnostics were calculated using satellite-derived and modelled velocity fields, providing multiple information on water mass transport and mixing (6) Finally, and for coral sites only, historical data of temperature were extracted (see (6) Historical data on coral sites) from satellite imagery to provide an historical overview of past heatwave experienced by the sampled coral reefs (since 2002 up to the sampling date).

**Table 3.** Data sets providing the provenance and the environmental context for future analysis and provided as raw measurements by sensors, from samples, and measurements aggregated at the sample, event and site levels.

| Name | Number of measurements | Variables | Link (final; see submission file for temporary tokens) |
| --- | --- | --- | --- |
| **Raw continuous measurements** |  |  |  |
| TSG | >590 000 | T, S | https://doi.pangaea.de/10.1594/PANGAEA.943675 |
| EDM | >15 000 | Aerosols concentration (0.25 – 32 µm) | https://doi.pangaea.de/10.1594/PANGAEA.943694<br>https://doi.pangaea.de/10.1594/PANGAEA.943691 |
| EIMS | >230 000 | O/Ar ratio | https://doi.pangaea.de/10.1594/PANGAEA.943714 |
| Optode | >280 000 | Oxygen concentration | https://doi.pangaea.de/10.1594/PANGAEA.943790 |
| Navigation | >1 271 000 | Navigation and Meteo | https://doi.pangaea.de/10.1594/PANGAEA.943725 |
| ACS | >411 000 | Chla, phytoplankton size, POC | https://zenodo.org/record/6449893 |
| BB3 | >350 000 | Backscattering, phytoplankton carbon | https://doi.pangaea.de/10.1594/PANGAEA.943793 |
| WSCD | >553 000 | relative DOM fluorescence (sd, n) | https://doi.pangaea.de/10.1594/PANGAEA.943739 |
| PAR | >830 000 | Photosynthetically active radiations (sd, n) | https://doi.pangaea.de/10.1594/PANGAEA.943740 |
| SMPS | >4600 | Aerosols concentration, particle size distribution (25 – 685 nm), sd | https://doi.pangaea.de/10.1594/PANGAEA.943856 |
| **Raw discrete measurements** |  |  |  |
| NUT | 849 | NO<sub>2</sub>, NO<sub>3</sub>, PO<sub>4</sub>, Si(OH)<sub>4</sub> | https://doi.pangaea.de/10.1594/PANGAEA.944289 |
| MTE-USC | 523 | Fe, Zn, Cd, Ni, Cu, Pb, Mn | https://doi.pangaea.de/10.1594/PANGAEA.944395 |
| CARB | 325 | Alkalinity, Carbonates, pH, pCO<sub>2</sub>, fCO<sub>2</sub>, HCO<sub>3</sub>, CO<sub>3</sub>, CO<sub>2</sub>, $\Omega_C$, $\Omega_{Aragonite}$ | https://doi.pangaea.de/10.1594/PANGAEA.944420 |
| FCM | 1041 | Pico-, Nano-, Picoplankton abundance and scattering | https://doi.pangaea.de/10.1594/PANGAEA.944490 |
| HPLC | 551 | Pigment concentrations | https://doi.pangaea.de/10.1594/PANGAEA.944281 |
| CTD | 4246 | T,S, conductivity, conductance, density, sound velocity, depth, pressure | https://doi.pangaea.de/10.1594/PANGAEA.943869 |
| **Environmental context at the granularity of sampling events** |  |  |  |
| Inline sensors + Almanach + Copernicus + Modis Aqua (2 and 12 pixels around) | 4155 | all Inline data with n, sd, quartiles, local sun/moon set/rize, local zenith, nutrients, hydrology, plancton quantities, Chla, PAR, PIC, POC, T, GSM, KD490 (with n, sd, and quartiles) | https://zenodo.org/record/6445609 |
| **Provenance metadata and environmental context at the granularity of samples** |  |  |  |
| Sample provenance | 57859 | georeference, sample unique identifier, logsheet links, environmental features and place names | https://zenodo.org/record/6299409 |
| All event level variables | 57859 | mean and std + dt, dx, dz from sampling timing, position and depth |  |
| **Environmental context at the granularity of sampling stations** |  |  |  |
| all event level variables | 655 | intercalibrated and combined version | https://zenodo.org/record/6474974 |
| Lagrangian Descriptors | 246 | Eulerian and Lagrangian diagnostics of water dynamic | https://zenodo.org/record/6453376 |
| **Environmental context at the granularity of coral sampling sites** |  |  |  |
| historical heat and cold stress indicators | 113 | TSA, DHW, recovery time etc... | https://zenodo.org/record/6499374 |
| raw time series | >6000 x 113 | SST at 1, 3 and 9 pixels, seasonal average, DHW, DCW |  |
| Reefcheck bleaching occurrence | 106 | Bleaching (% of colony or % of population) | https://zenodo.org/record/6511406 |
| **Photo annotations** |  |  |  |
| Qualitative photographic annotations | 5606 photo, 2216 colonies | identification to the genus level, algal contact (genus of algae), presence of boring organisms (type), contact with sediment, presence of predation marks | https://zenodo.org/record/6364768 |
| Taxonomic annotations of coral diversity (CDIV) surveys | 2470 | 18S based taxonomic annotations, corresponding morphological annotation based on photo | https://zenodo.org/record/6327048<br>https://ecotaxa.obs-vlfr.fr/prj/4176 |

### (1) Raw measurements from samples or sensors

From sensors, the measurements were standardized at the minute scale when possible (including standard deviation and the number of observations within the minute when available) and accompanied with their UTC time and GPS position. These data sets regroup data obtained from the [TSG] the [ACS] the [WSCD] the [BB3] the [EIMS] the [optode], the [EDM], the [SMPS], the [PAR] and the navigation data. These are available as ten distinct data sets, one for each package of sensors. Similarly, measurements made from discrete samples collected on board Tara (see Methods Section 3.3), together with quality assessment flags, are provided as six distinct data sets, one for each type of analysis ([NUT], [MTE-USC], [CARB], [FCM], [HPLC], and [CTD]). For [CARB], additional parameters of the carbonate system were calculated with CO2SYS.m v3.1.1^31^ using in situ temperature, total alkalinity, total dissolved inorganic carbon, salinity, phosphate and silicate concentrations as inputs together with recommended parameters^32–35^ (K1K2=10; KSO4=3; KF=2; BOR=2). Data sets are available in open access at the Data Publisher for Earth & Environmental Science PANGAEA.

### (2) combined version at the event level

A compilation of all environmental measures obtained during a given sampling event was produced by compiling the boat’s sensor data available during the time-lapse of the station and measurements originating from satellite imagery (MODIS-AQUA satellite - Level 3 mapped product, 8-day average, 4km resolution) recovered using OpenDAB protocols at https://oceandata.sci.gsfc.nasa.gov. The zone corresponding to the station position and date was recovered either by taking a two-pixel buffer around the given location (total zone being a 5 by 5 pixels square of 20 km side) and in order to propose an alternative measure in the inevitable case where clouds were present an alternative 12-pixels buffer was taken (total zone being a 25 by 25 pixels square of 100 km side).

The corresponding variables recovered are chlorophyll *a*^36^ (OCx algorithm^37^, [Chl_Sat]; mg m^-3^), the sea surface temperature^38^ (4µm shortwave algorithm; [T_Sat]; °C), daily mean photosynthetically available radiation at the ocean surface^39^ ([PAR_Sat]; Einstein m^-2^ d^-1^), concentration of particulate inorganic carbon^40^ ([PIC_Sat]; mol m^-3^), concentration of particulate organic carbon^41^ ([POC_Sat]; mol m^-3^), the diffuse attenuation coefficient for downwelling irradiance at 490 nm^42^ ([Kd490_Sat] related to light penetration in water column modified by particulate matter; m^-1^), and the particulate backscattering coefficient at 443 nm derived from the Garver-Siegel-Maritorena algorithm^43^ ([GSM_Sat] which gives a good indication of the concentration of suspended organic and inorganic particles such as sediments in the water; m^-1^).

This compilation of environmental data at the scale of the event was further enriched using data from reanalyzed (ie. forced with observations) operational models obtained from Copernicus Marine Services (GLOBAL_REANALYSIS_PHY_001_030^44^, daily mean for sea surface height, salinity, temperature, current speeds, mixed layer depth; GLOBAL_REANALYSIS_BIO_001_029^45^ daily mean for Chl a, phytoplankton carbon, O2, NO3, PO4, SIOH, Fe concentrations, Primary production, pH and CO2 partial pressure and GLOBAL_REANALYSIS_WAV_001_032-TDS^46^ for sea surface waves) but also using almanach^47, 48^ to calculate essential sun and moon parameters (position, rises and sets, phase, etc).

### (3) Environmental context at the granularity of samples

The environmental context of all samples collected during the Tara Pacific Expedition is available together with the provenance file in open access at Zenodo. The environmental context of each sample is provided based on environmental data sets described above for continuous and discrete measurements, as well as those generated from almanacs, satellite imagery and models.

Environmental context is provided in eleven UTF-8 encoded tab-separated-values files, all with the same structure, but each providing a different statistic: number of values (n), mean value (mean), standard deviation (stdev), 05, 25, 50, 75 and 95 percentiles (P05, P25, P50, P75, P95), lag in time (dt), i.e. difference between the collection date/time of the sample and that of the environmental context provided, lag in horizontal space (dxy), i.e. distance between the collection location of the sample and that of the environmental context provided, and lag in vertical space (dz), i.e. difference between the collection depth/altitude of the sample and that of the environmental context provided. Missing value terms are: “nav” = not-available, i.e. the expected information is not given because it has not been collected or generated; “npr” = not-provided, i.e. the expected information has been collected or generated but it is not given, i.e. a value may be available in a later version or may be obtained by contacting the data providers; “nac” = confidential, i.e. the expected information has been collected or generated but is not available openly because of privacy concerns; “nap” = not- applicable, i.e. no information is expected for this combination of parameter, environment and/or method, e.g. depth below seabed cannot be informed for a sample collected in the water or the atmosphere

### (4) Simplified version at site level

In some cases, certain parameters were not available at specific sampling sites due to technical issues or sensor availability, however, various basin scale studies and statistical tests require a complete dataset for all sampled sites. During the Tara Pacific expedition, many parameters were concurrently measured in-situ, estimated from remote sensing and/or modelled. For instance, sea surface temperature was measured on the boat using the thermosalinograph included in the underway system, but also with satellite and estimated from a model. Each of these three modes of acquisition have their caveat and accuracy, however, within a certain confidence interval, missing in-situ data can be replaced by its remotely sensed or modelled equivalent. We provide here a simplified version at the sampling site level by replacing missing in-situ data by their closest and most accurate satellite or modelled equivalent. In each case, in-situ data was considered as the most accurate source of data, with a preference to HPLC pigments analysis followed by measurements done by the ACS, while satellite and modelled data were used only if in-situ data was not available. We evaluated the accuracy of ACS and of each satellite and modelled datasets by linear regressions with their in-situ counterparts. A bias of the modelled or satellite data was identified when the slope of the regression was different to 1 and/or an intercept was different to 0. The satellite and modelled data were forced to match the in-situ data by dividing by the slope and subtracting the intercept. This is the case for SST. When large bias persisted between matchups with observations, the corrected data was not used to replace missing in-situ data. This is the case for chl. The same approach was then applied to fill missing data with modelled values (MERCATOR-Copernicus).

A correction for the bias in the following variable was applied for SST, SSS, PO4, and SiOH. As previously done, if large bias persisted between observations and corrected data, they were not used to replace missing in-situ data. This is the case for chl, NO3, and Fe.

The [MTE] samples were sometimes sampled in the afternoon instead of the morning alongside all the other water samples, thus were located in between two sampling stations. These [MTE] samples could not be assigned to a sampling station following the criterion presented in the section 3, therefore, the missing values of the corresponding morning stations were interpolated linearly.

The same approach was used for pH measurements, with a preference from measurements provided by total carbonate system quantifications, followed by direct pH measurements and then modeled values (MERCATOR-Copernicus).

### (5) Lagrangian and Eulerian diagnostics

In order to provide a description of the dynamical properties of the water masses sampled, different Eulerian and Lagrangian diagnostics were calculated. Here, we report a general description of the information each of them provides. In the next subsection, we provide the details of how they were calculated for each station.

The following Eulerian diagnostics were calculated: Absolute velocity ([Uabs], m s^-1^): sqrt(u^2^+v^2^), where u and v are the zonal and meridional components of the horizontal velocity field used (described below); Kinetic energy ([Ekin], m^2^ .s^-2^): 0.5*(u^2^+v^2^); Divergence ([EulerDiverg], d^-1^): du/dx + dv/dy; Vorticity ([Vorticity], d^-1^): dv/dx - du/dy; Okubo-Weiss ([OW], d^-2^): s^2^- vorticity^2^, where s^2^ is (du/dx-dv/dy)^2^ + (dv/dx+du/dy)^2^. If negative, it indicates that the station sampled was inside an eddy.

The following Lagrangian diagnostics were calculated: Finite-Time Lyapunov Exponents ([Ftle], d^-1^, Shadden et al., 2005): it indicates the rate of horizontal stirring, and it is a means to quantify the intensity of turbulence in a given region. FTLE are commonly used to identify Lagrangian Coherent Structures, i.e. barriers to transport. In this case, a strong FTLE value indicates a region separating water masses which were far away backward in time. Lagrangian betweenness^49^ ([betw], adimensional): this diagnostic draws inspiration from Lagrangian Flow Network Theory^50^. It can identify regions which act as bottlenecks for the circulation, in that they receive waters coming from different origins, and that are then spread over several different destinations. These can represent possible hotspots driving biodiversity^49^. Lagrangian Divergence^51^ ([LagrDiverg], d^-1^). This diagnostic was calculated by integrating the Eulerian divergence along the backward trajectories. If positive, it indicates a water mass that, during the previous days, was subjected to a strong divergence, thus to a possible upwelling. If negative, it indicates a strong convergence, thus possible downwelling. Retention Time^52^ ([RetentionTime], d). This diagnostic indicates how many days a water mass has spent inside an eddy in the previous period. If the water mass is outside an eddy, then its retention time is set to zero.

#### (5.1) Extraction of the Eulerian and Lagrangian diagnostics

For each of the 246 stations sampled, we proceeded as follows.

We identified the water mass sampled at the given station. This was considered as a stadium shape with the two semi-circles centered on the starting and ending points of the transect, respectively. The radius of the stadium semi-circles was considered 0.1°, which is in accordance with previous studies^49, 53, 54^. The stadium was filled with virtual particles separated by 0.01°.

For each virtual particle inside the stadium shape, we calculated a Eulerian or Lagrangian diagnostic (described above). The Eulerian diagnostics were extracted directly from the velocity field of the day of sampling. Concerning the Lagrangian diagnostics, these were obtained by advecting the virtual particle backward in time for an amount of time τ from the day of sampling day_S. For the Lagrangian betweenness, the advection was performed between day_S+τ/2 and day_S-τ/2, so that the advective time window was centered on the sampling day (details in^49^).

For the Lagrangian diagnostics, we used the following advective times τ: 5, 10, 15, 20, 30, and 60 days. The only exception is the retention time, which, by construction, was calculated only with the largest advective time, namely τ=60 days. Once that, a given diagnostic (Eulerian or Lagrangian) was calculated for all the virtual particles filling the stadium shape, we calculated the mean value, and the 25, 50, and 75 percentiles. The percentiles were calculated in order to quantify the spatial variation of the diagnostic inside the stadium shape. Therefore, we associated each station with four values (mean, 25, 50, and 75 percentiles) of a given diagnostic.

Furthermore, two different velocity fields were used, which are described as follows.

#### (5.2) Velocity fields and trajectory calculation

Both the velocity fields were downloaded from E.U. Copernicus Marine Environment Monitoring Service (CMEMS, http://marine.copernicus.eu/). The first velocity field used was MULTIOBS_GLO_PHY_REP_015_004^55^ [GlobEkmanDt]. This was produced by combining the altimetry derived geostrophic velocities and modelled Ekman surface currents. It had a spatial resolution of 0.25° and a temporal resolution of one day. The second velocity field was GLOBAL_REANALYSIS_PHY_001_030^44^ [GloryS12]. It was obtained by a NEMO model assimilating altimetry and other observations. It had a spatial resolution of 1/12° and a temporal resolution of 1 day.

### (6) Historical climate data and indices for climate variability for coral collection sites

It’s becoming increasingly clear that stress resilience, in particular thermal tolerance, is shaped not only by maximum monthly mean temperatures (MMMs), but also by long-term and short-term climate variability, even at the scale of reefs^56–58^. In order to provide an overview of past climate variability and heatwaves experienced by corals sampled at each site, we built a high-resolution historical dataset that spans from 2002 to each sites’ sampling date. Ocean skin temperature (11 and 12 µm spectral bands longwave algorithm) was extracted from 1km resolution level-2 MODIS-Aqua and MODIS-Terra from 2002 to the sampling date and from level-2 VIIRS-SNPP from 2012 to the sampling date. Day and night overpasses were used to maximize data recovery. Following recommendations from NASA Ocean Color (OB.DAAC), only SST products of quality 0 and 1 were used. The 9 closest pixels to the sampling sites of each scene were extracted. All the extracted pixels from the 3 platforms were then averaged daily to obtain daily SST averages and standard deviations time series for each sampling site, from 2002 to the sampling date.

Each time series was first averaged on a Julian day basis to provide a seasonal average. This yearly seasonal average was triplicated and concatenated into a 3-year seasonal cycle to apply a digital low pass filter on the middle year without generating artifacts. A digital low pass filter (filter order 3, pass band ripple 0.1; “filfilt” function in matlab) with 36 Julian days windows was applied to the concatenated time series to remove high frequency noise. The middle year was then extracted from the concatenated time series to recover the seasonal cycle. The sea surface temperature anomaly was calculated as the SST minus the seasonal cycle over the full time series. Considering the short periods of missing data (mean of the 95th percentile of the duration of consecutive days with missing data: 9.8 ± 4.1 days), the missing values in the SST and SST anomaly time series were linearly interpolated in order to calculate thermal stress indices. The SST anomaly frequency was calculated as the number of days over the past 52 weeks when the SST anomaly is greater than or equal to 1 °C. Thermal stress indices relevant to coral reef health were then calculated using methodology developed for the Coral Reef Temperature Anomaly Database (CoRTAD) data base^58^ (Table 4). Events of cold temperature accumulation were also reported to cause bleaching and mortality^59, 60^, therefore, the same set of indices were calculated for cold stress adapting the CoRTAD method, but using the minimum weekly climatologies (Table 4). Further to that, we checked for previous occurrences of bleaching events at sampled reef sites by matching island coordinates to the Reef Check dataset (reefcheck.org) obtained from Sully et al 2019^56, 61^. For each Tara Pacific island, coordinate we determined that Reef Check site that was closest (in terms of distance in km) and considered only Reef Check data that was within a 10 km circumference.

**Table 4:** Description of historical SST values and thermal stress indices calculated following CoRTAD^58^ method and modified to also represent cooling events.

| Name | Acronym | Description | Reference |
| --- | --- | --- | --- |
| SST daily average 9 pixels | [sst_mean_9pixel] | Daily average of the 9 closest pixels around the sampling site |  |
| Seasonal average 9 pixels | [seasonal_average_9pixel] | Seasonal average SST calculated from 2002 to the sampling date |  |
| SST anomaly 9 pixels | [SST_anomaly_9pixel] | SST anomaly calculated as: sst_mean_9pixel minus seasonal_average_9pixel |  |
| SST daily average interpolated | [SST_mean_interpl] | SST daily average with missing values interpolated linearly |  |
| SST anomaly interpolated | [SST_anomaly_interpl] | SST anomaly with missing values interpolated linearly |  |
| SST anomaly frequency | [SST_anomaly_freq] | number of days in the past 52 weeks when SST_anomaly_interpl $\geq 1$ °C | CoRTAD |
| Heat Thermal Stress Anomaly | [TSA_heat] | Daily SST average interpolated minus the maximum weekly climatology | CoRTAD |
| TSA heat frequency | [TSA_heat_freq] | number of days in the past 52 weeks when TSA_heat $\geq 1$ °C | CoRTAD |
| TSA degree heating week | [TSA_DHW] | sum of the past 12 weeks when TSA_heat is greater than or equal to 1 °C | CoRTAD |
| TSA degree heating week frequency | [TSA_DHW_freq] | number of days in the past 52 weeks when TSA_DHW is greater than or equal to 1 °C | CoRTAD |
| Cold Thermal Stress Anomaly | [TSA_cold] | Daily SST average interpolated minus the minimum weekly climatology | Custom made |
| TSA cold frequency | [TSA_cold_freq] | number of days in the past 52 weeks when TSA_cold $\leq -1$ °C | Custom made |
| TSA degree cooling week | [TSA_DCW] | sum of the past 12 weeks when TSA_cold is lower than or equal to -1 °C | Custom made |
| TSA degree cooling week frequency | [TSA_DCW_freq] | number of days in the past 52 weeks when TSA_DCW is lower than or equal to 1 °C | Custom made |

A condensed table containing single values associated with each sampling site was created extracting the minimum, maximum, sum, averages, standard deviations, and value recorded at the sampling day of each of these indices (detailed in the readme file provided with the dataset). Additional metrics of the last heating and cooling events as well as the time of recovery is also provided to represent the state of thermal stress at the day of sampling.

### (7) Coral photographic resources and annotations

The [PHOTO] resource consists of two datasets. The first, obtained from the [SCUBA-3X10] protocol, was annotated for genus validation, gross morphological characteristics of the colony, algal contact, presence of boring organisms, sediment contact, predation, and health factors (such as presence of disease and coloration). The acquisition protocol of these annotations is described below. This dataset is also used for the description of morphotypes within each genus for taxonomic annotation in combination with genetic data. The second dataset, obtained following [SCUBA- SURVEY] protocol was used for the taxonomic annotation (as close to genus level as possible) of the coral host of the [CDIV] samples. Of a total of 2,470 CDIV samples, 1711 samples had one or more pictures associated (3,085 total pictures), 759 samples had no photos. Overall, 11,460 coral photographs were generated and annotated allowing for a permanent record of all colonies sampled. All [PHOTO] were transferred to EcoTaxa^62^.

#### (1)#Manual Annotations of *in situ* colony (CO) photos

Photo analysis for the genus validation and environmental context was conducted using Matlab with code developed and written specifically for the Tara Pacific Expedition^63^. Photos were annotated individually, and annotations were conducted from January to April 2020. To prevent observer bias, photos were randomized, and the annotator was blind to any information regarding the location or the sampling site. The analysis included 1) identification to the genus level, 2) algal contact with types of algal genus if identifiable (Halimeda, Turbinaria, Dictyota, Lobophora, Crustose Coraline Algae (CCA), Sargassum, Galaxaura, other), 3) presence of boring organisms with types if identifiable (Bivalve, Spirobranchus, Tridacna, Urchin, Other Polychaete, Sponge, and Other), 5) contact with sediment (sand), 6) presence of predation marks. Most annotations were boolean operators (yes/no) with identifications added if possible. Several indicators of coral health were also annotated such as if the coral looked unhealthy or showed tissue loss (Yes/No), coloration (light, normal, dark, or bleached), and presence of a pigmentation response (Yes/No). If a pigmentation response was present, the annotator was prompted to determine if it was trematodiasis (Yes/No). Finally, additional notes included but were not limited to the quality of the photo (blurry, poor visibility, coloration), contact with neighbouring hard or soft coral colonies, fish presence in the photograph, snail(s), or hermit crab(s) on the coral, an object in the photograph, etc.

#### (2)#Taxonomic annotations of coral diversity (CDIV) surveys

All images imported in EcoTaxa have been identified at the genus level by taxonomic experts, and crosslinked with genomic identification from metabarcoding based on the V9 region of the 18S rDNA. Analysis of the 18S marker aimed to generate coral host taxonomic annotations to the level of genus for every sample. The annotation was generated based on each sample’s most abundant 18S sequence by aligning to the NCBI ‘nt’ database with taxonomic labels. A ‘lowest common ancestor’ approach was used when there were multiple best hits. These alignment-based annotations were verified phylogenetically (i.e. taxonomic similarity agreed with sequence similarity). More than half of the samples were not annotated at genus or better level using this approach, due to the lack of resolution of the 18S V9 marker. Where available, host taxonomic assignments were based on photo annotations. Otherwise, 18S-based annotations were used.

## Technical Validation

Numerous steps of quality control were operated at different levels of acquisition to ensure good quality of the different datasets and may vary depending on the type of measurement operated and if it originates from sensors on-board or from samples.

## Inline measurements, models and satellite data validity

[PAR] measurement validity was checked by first removing physically wrong data (ie. values greater than 0.45 μE/cm2/sec or lower than 0 μE/cm2/sec) and compared with clear sky matchup measurements from MODIS-Aqua & Terra. Comparison confirmed the good agreement between datasets but also the absence of sensor drift. Temperature and salinity were acquired by the [TSG]. The quality of the whole time series was manually checked, and the temperature validity was assessed by comparing the temperature reading of the two sensors placed at two different places along the inline system. Potential drifts of the temperature sensor was investigated by comparing the temperature time series with satellites’ sea surface temperature. Salinity measurements where intercalibrated against unfiltered seawater samples [SAL] taken every week from the surface ocean, and corrected for any observed bias. Moreover, temperature and salinity measurements were validated against Argo floats data collocated with Tara. The [ACS] absorption and attenuation signal due to dissolved matter, drift, and biofouling were estimated between two filter events by interpolating filtered water absorption and attenuation following the shape of the [fdom] from the [WSCD], when available. This method improves data quality in case of strong variation of dissolved matter absorption that the frequency of filter event would not capture properly (e.g. approaching coastal waters or entering a lagoon). When [fdom] data was not available, the filtered absorption and attenuation were linearly interpolated between filter events before being remove from the total absorption and attenuation. From November 13, 2016 to May 6, 2017, the [BB3] was located upstream of the switch system, thus measured total (non-filtered) water all the time. During this period, the volume scattering coefficient of seawater was removed from the raw data counts to obtain the particulate backscattering coefficient [bbp]. The biofouling and instrument drift were estimated comparing values before and after each cleaning events. The biofouling was estimated between two cleaning events by fitting an exponential or linear model to the raw data before removing it from the signal. We advocate to use this period with caution as the data was corrected with theoretical assumptions (i.e. pure seawater scattering and linear or exponential biofouling) that may differ from reality. From May 7^th^ 2017 to the end of the expedition, the [BB3] was located downstream of the filter-switch system so that, like for the [ACS] processing, the biofouling signal could be estimated and removed between two filter events and [bbp] quality improved. The correspondence between total particulate scattering [bp] estimated from the [ACS] and [bbp] was investigated for the whole expedition. [bbp] values were discarded when [bbp]/[bp] was unusually low (< 0.002; see range of [bbp]/[bp] in natural waters^64^). A similar modelling and correction for biofouling than the one performed for the [BB3] was applied to the [WSCD] data. The [PAR], [TSG], [BB3], [ACS], and [WSCD] data were processed following the last recommendations for processing inline^23^, using custom software available at https://github.com/OceanOptics/InLineAnalysis. The entire time series of measurement were automatically QC to remove artifacts and manually checked and QC for obviously inaccurate measurements due to saturated sensor, low flow rate, bubbles, or poor filtered seawater measurements. The full processing and QC procedure and reports could be accessed together with each dataset.

## Sample measurements technical validation

For nutrients [NUT] samples a quality check was done in several steps. First a visual inspection was done to determine if samples were overfilled or not frozen vertically which may induce sample leakage during the frosting procedure. Secondly any readings too close to detection limits or when duplicate measurements differed by more than 10% were flagged. In this last case, when the difference between two values of the same sample is greater than 10%, it is considered that the high value is not acceptable and is not reported. Finally, the overall quality of the dataset was established by comparing measurements values with Copernicus Marine Services modelling outputs.

For trace metals ([MTE-USC]), any samples in which concentrations were close to detection limits were flagged. A standard produced by the GEOTRACES program (coastal surface seawater standard) was included in each sample run. If the metal concentrations of the standard were outside the GEOTRACES community consensus values, the sample run was rejected. Trace metal concentrations had an average error of 5%.

[HPLC] samples were analysed as described in Ras et al 2008. All pigments peaks were inspected and quality controlled as good, acceptable or qualitative. Any measurements below detection limits were disregarded.

[FCM] samples were analysed with a FACS Canto II Flow Cytometer equipped with a 488 nm laser^32^ and every measurement where cell populations were either complicated, needed manual curation or were impossible were flagged.

## Nets collection validity

To estimate the technical validity of the different nets collection we analysed the raw abundance of living organisms collected conjointly by the [HSN-NET-300] and [MANTA-NET-300] at the same stations, but sequentially in time. Indeed [MANTA-NET-300] is operated at different speeds (3 knots maximum) compared to [HSN-NET-300] (9 knots maximum) and therefore were not deployed simultaneously. Manta nets are commonly used and recognized as a reference type of net while investigating surface plankton^65–67^ and we therefore used a set of 24 stations where both were deployed concurrently to estimate the efficiency of the [HSN-NET-300]. For this [F300] samples collected by both nets were imaged using the ZooScan^68^ to obtain images of each object collected. Images were then transferred to EcoTaxa^62^ and sorted taxonomically to the deepest taxonomic level possible. All results were used to calculate the normalized biovolume size spectra^69^ (NBSS) of living organisms for both nets, which is an analogue to abundance per size categories. This NBSS spectra allows investigating the potential under- or over-sampling while investigating it over various sizes of organisms. The NBSS of both nets were giving about the same order of magnitudes of abundances (Figure 4A) and when inspected along the size spectra between pairs of observations (Figure 4B) they did not differ largely from 1:1 in 13 cases over the different deployments. A large variability between nets could however be observed at a few stations which could possibly be caused by local plankton patchiness^70^ resulting in more variability for [HSN-NET-300] and less for [MANTA-NET-300] due to larger sampling volume. Overall, we can conclude that [HSN- NET-300] and [MANTA-NET-300] are collecting plankton with a relatively similar efficiency even if the larger sampling volume of [MANTA-NET-300] allows a better collection of larger, rare, organisms, as seen from spectra extending to larger sizes (Figure 4A). Nevertheless, these results show that the use of [HSN-NET-300] may be really useful for underway zooplankton sampling in the situations when it is not possible to stop the ship for regular sampling or on ships of opportunity.

**Figure 4:**
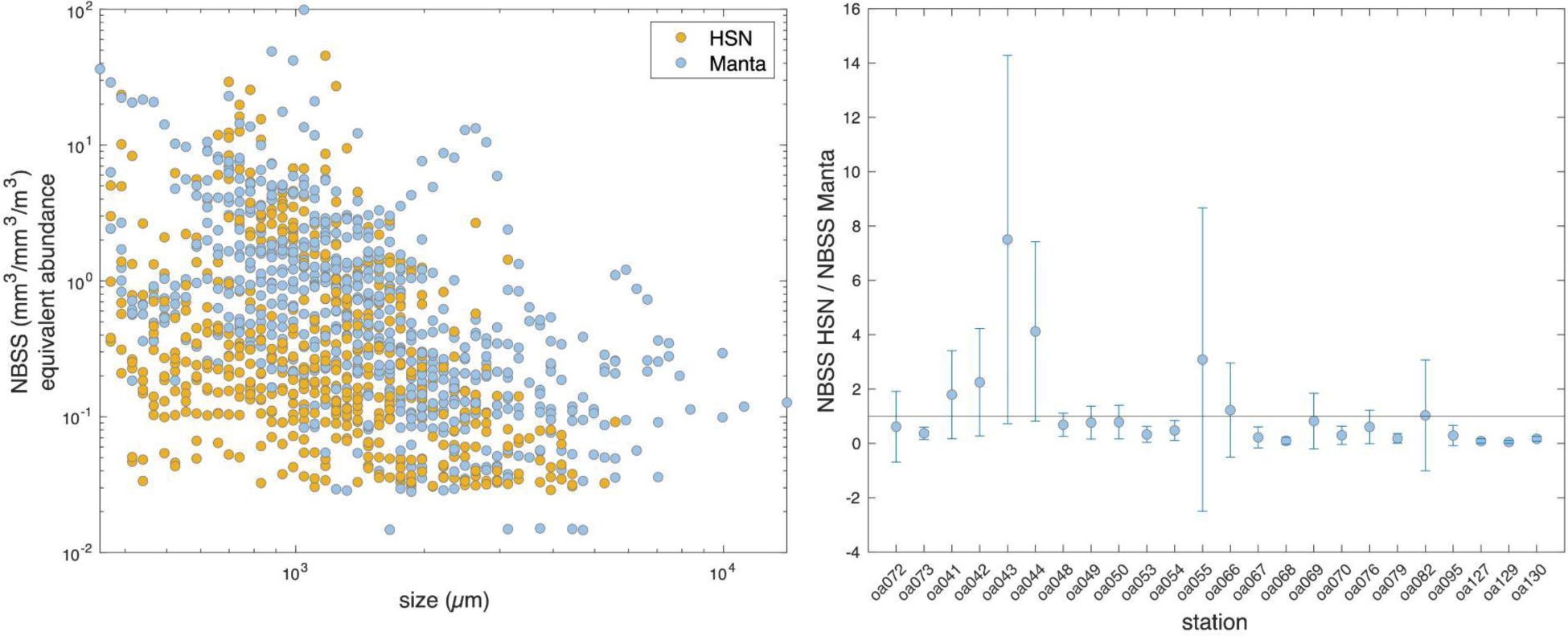
Technical validation of net sampling. Comparison of normalized biovolume size spectra (NBSS) of living organisms sampled with [HSN-NET-300] and [MANTA-NET-300] over a set of 24 stations where both were deployed together. From both NBSS, a sampling efficiency of the HSN net compared to the MANTA net was calculated as a mean and standard deviation over all the size classes considered.

## Overall biogeochemical data validity

To assess the overall quality and homogeneity of the collected environmental parameters, we conducted a quick multivariate exploration of the dataset to compare it with known biogeography of biogeochemical provinces^71, 72^ and their associated biogeochemical signatures. For this, we first used data simplified at the site version (see section 4 of Data records), selected only datasets providing a full overview over the geographical range of the expedition, used a box-cox transformation and centred-reduced each variable to equally consider those. This dataset was then analysed through a PCA analysis (Figure 5). The 3 first components of the PCA analysis were recovered to code for a RGB (red, green, blue) color-coding of each station and better visualize the biogeochemical signature of the station on a map. Finally, those were compared with known biogeochemical provinces extracted from^72^. Despite the different temporal resolution between instantaneous sampling and biogeochemical provinces representing a consensus over several years and seasons, we can see that the main biogeochemical provinces (and associated macroscale oceanic features) as well as their progressive boundaries are well captured by our sampling scheme. Among the notable features, the western Pacific coast of Americas are marked by a strong upwelling signature (with high amount of nutrients and trace metals), the southern Pacific gyre with a high salinity but a low iron and silicate concentration, the central Pacific zone is characterized by high temperature, light and sea surface height, small phytoplankton size (high gamma), with low chlorophyll a and low NO3 and trace metals (Ni, Cu, Zn, Pb or Cu) concentrations, with the exception of the few stations centred on the equator which clearly display some indicators of local upwelling such as those potentially created by the equatorial upwelling. This first overview clearly shows correspondence with known features related to nutrients and nutrient limitation of plankton, trace metals or even global biogeochemistry^73–75^ and further shows that the sampling scheme used allowed to sample corals and plankton across a large variety of environmental constraints either on oceanographic, climatic or chemical aspects. The same analysis repeated only using sites realized around islands further confirms this large variety of environmental constraints (Figure 6). To evaluate the variety of the past temperature history, and notably the impact of past seasonality and heat/cold waves, we further reproduced this analysis using historical temperature and heat/cold waves experienced on coral sites. However, since temperature anomalies and their accumulated degree cooling weeks (DCW) could be negative, only a basic normalization of data was made since box-cox normalization is not suited for negative values. The first axis of the PCA separate islands that suffered intense and recurrent heat-waves (positive values) from those that rather experienced cold-waves (negative values) while the second axis separate cold and highly seasonal islands (positive values) from islands with warm environments with low seasonality (negative values). This analysis further confirms that the selected location also displays a full variety of past history of temperature and heat-waves but also reflects known geographical patterns of bleaching events^56, 76^.

**Figure 5:**
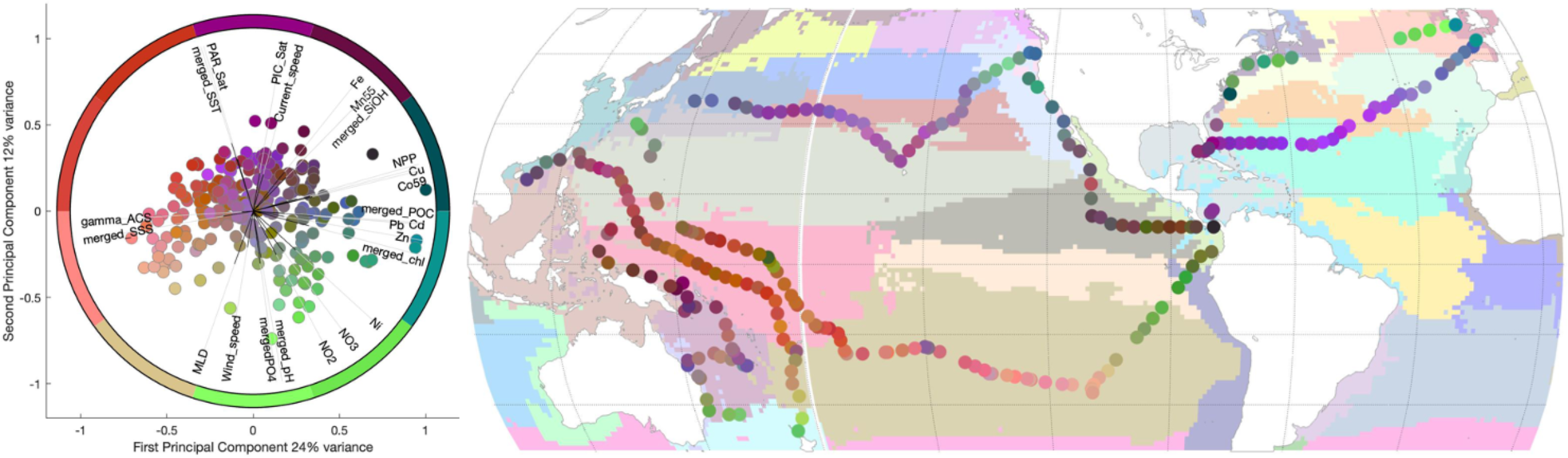
Technical validation of the main hydrological and biogeochemical environmental variables compared with biogeochemical provinces as extracted from^72^. Environmental data were normalized through a box-cox normalization and analysed through a PCA analysis to better display their typical environmental signature. The position of each station in the 3 first axes of the PCA were further used to to provide a red blue green color-coding, allowing to project their environmental signature on a map and compare it with known biogeochemical provinces.

**Figure 6:**
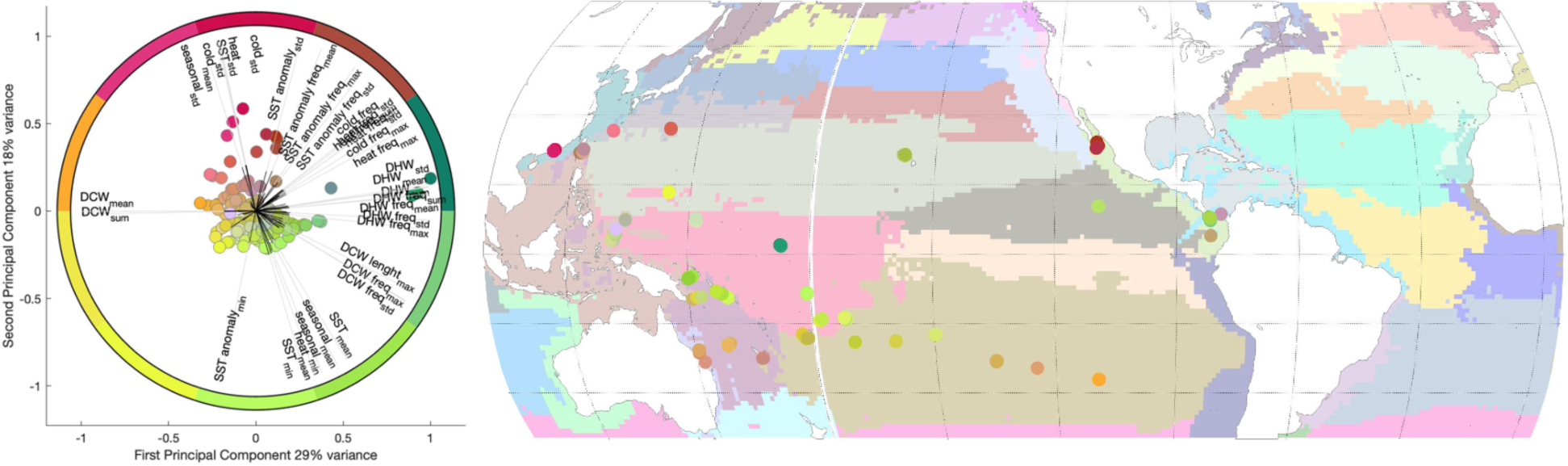
Technical validation of the historical SST heatwaves and cold waves parameters compared with biogeochemical provinces as extracted from^72^. Environmental data were normalized and analysed through a PCA analysis to better display their typical environmental signature. The position of each station in the 3 first axes of the PCA were further used to provide a red blue green color-coding, allowing to project their environmental signature on a map and compare it with known biogeochemical provinces.

## Usage Notes

We recommend paying close attention to the various quality flags provided with the raw datasets to avoid using lower quality data if needed. Similarly, to provide the more complete set of observations for each sample, we provided the lag in time (dt), as well as distance in horizontal (dxy) and vertical (dz) space, between the collection timing, latitude/longitude and depth/altitude of the sample and that of the environmental context provided. Depending on the scientific question, future users are encouraged to carefully define reasonable time lag and distances to consider in their study, to avoid including unrealistic associations between samples. Moreover, we extracted contextual data at the event level to simplify the data extraction task. We also provide simplified version at the site level by combining and cross-calibrating all similar variables (e.g. using different sources of SST data to fill gaps of missing data and obtain one merged SST variable). We prioritised observations originating from in-situ samples over satellite data, and over modelled data (MERCATOR), and evaluated their correspondence by linear regressions. Potential biases of satellite and modelled data in comparison to in-situ data were corrected applying the slope and intercept of their linear regression to force satellite and modelled data to best match in-situ data. Similarly, we also chose to interpolate some environmental variables that were sampled only few hours before or after the site itself to maximize data recovery for each sampling station. We acknowledge merging different sources of data can introduce differences in variance depending on the source of data used, therefore, we encourage the user to cautiously evaluate the relevance of this merged dataset for their study. Considering the intrinsic heterogeneity of variance between the different datasets, and their potential non-normal distribution, we recommend using appropriate normalisation methods before any multivariate statistical analysis. Here we chose to use box-cox transformation and centred-reduced each variable.

In this version of the dataset the satellite data used is 8-days averages while the in-situ measurements are instantaneous measurements of optical properties averaged over the station sampling period. The 8-days averaging tend to attenuate extreme values and reduce the potential differences between stations. While suited for macro-ecological processes which depends on large temporal and spatial variations of their environment, the use of 8-day averages satellite products could be inaccurate to study shorter life cycles of the pico-, nano and micro-plankton.

Moreover, phytoplankton can adjust their light harvesting pigment concentrations according to light exposure, nutrient availability and temperature. These variations are negligible over periods shorter than a day but can become significant over 3-5 days^77^ and references therein. Therefore, we advise the users to cautiously use the merged bio-optical variables of this dataset and to verify its compatibility with the research question and potentially replace this 8-day average with shorter time observations if available. As presented in section “3.3. Continuous measurements”, the [poc] was estimated from the underway system, both using the measured [cp]^27^, and [bbp]^28^. The [BB3] sensor have a low signal-to-noise ratio due to its high sensitivity to bubbles in the water line and to accumulation of particles in the sensor, therefore, the [poc] estimated from the [BB3] was used to fill the missing [poc] estimated from the [ACS]. When the [bbp] from the [BB3] was used to estimate [POC], the [bbp] values from the 470 nm wavelength were prioritized over the 532 nm wavelength and 650 nm wavelength and the same merging method was applied to correct for bias between [poc] estimated from the [ACS] and the [BB3], and between wavelength of the [BB3].

## Code Availability

The different codes used to process the different datasets are indicated within the text and are repeated here and includes:

– Inline optical processing (https://github.com/OceanOptics/InLineAnalysis)
– Satellite products used^36, 38–43^
– Mercator products^44–46, 55^ used.
– Astronomical almanac to calculate sun/moon position and day-nights parameters from sites positions and time^47, 48^.
– Additional parameters of the carbonate system were calculated with CO2SYS.m v3.1.1^31^ using in situ temperature, total alkalinity, total dissolved inorganic carbon, salinity, phosphate and silicate concentrations as inputs together with recommended parameters^32–35^ (K1K2=10; KSO4=3; KF=2; BOR=2).
– Ecotaxa^62^ server github (https://github.com/ecotaxa/ecotaxa).
– EcoTaxa data processing (https://github.com/ecotaxa/ecotaxatoolbox)
– Morphological qualitative annotations^63^.

## Acknowledgements

We are keen to thank the commitment of the people and the following institutions for their financial and scientific support that made this singular expedition possible: CNRS, PSL, CSM, EPHE, Genoscope/CEA, Inserm, Université Côte d’Azur, ANR, agnès b., UNESCO-IOC, the Veolia Environment Foundation, Région Bretagne, Billerudkorsnas, Amerisource Bergen Company, Lorient Agglomeration, Smilewave, Oceans by Disney, the Prince Albert II de Monaco Foundation, L’Oréal, Biotherm, France Collectivités, Fonds Français pour l’Environnement Mondial (FFEM), the Ministère des Affaires Européennes et Etrangères, the Museum National d’Histoire Naturelle, Etienne BOURGOIS, the Tara Ocean Foundation’s teams and crew.Tara Pacific would not exist without the continuous support of the participating institutes. The authors also particularly thank Serge Planes, Denis Allemand and the Tara Pacific consortium. This study has been conducted using E.U. Copernicus Marine Service Information and Mercator Ocean products. We acknowledge funding from the Investissement d’avenir project France Génomique (ANR-10-INBS-09). FL is supported by Sorbonne Université, Institut Universitaire de France and the Fondation CA-PCA. The in-line and atmospheric optics dataset was collected and analysed with support from NASA Ocean Biology and Biogeochemistry program under grants NNX13AE58G and NNX15AC08G and the HPLC processing under NSF award 2025402 to University of Maine. JMF is supported by a research grant from Scott Eric Jordan. NCo was supported by a grant from the Simons Foundation/SFARI (544236). Work from MC, RMM, ER, DF, PF, EG is supported by the French Government (National Research Agency, ANR) through the grant “Coralgene” ANR-17-CE02-0020 as well as the “Investments for the Future” programs LABEX SIGNALIFE ANR-11-LABX-0028 and IDEX UCAJedi ANR-15-IDEX-01. NCa and YL were supported by the “Laboratoire d’Excellence” LabexMER (ANR-10-LABX-19) and co-funded by a grant from the French government under the program “Investissements d’Avenir”. FL, SP, CdV and ZM are funded by the European Union’s Horizon 2020 research and innovation programme “Atlantic Ecosystems Assessment, Forecasting and Sustainability” (AtlantECO) under grant agreement No 862923. The support of Pr. Alan Fuchs, President of CNRS, was crucial for the success of the surface sampling undertaken during the *Tara* Pacific expedition. We thank A. Gavilli from TECA Inc. France, and E. Tanguy and D. Delhommeau from the Institut de la Mer, Villefranche-sur-Mer for the helpful collaboration in the conception of the High Speed Net and the Dolphin systems. The AT and CT data were analysed at the SNAPO-CO2 service facility at LOCEAN laboratory and supported by CNRS-INSU and OSU Ecce-Terra. We thank the EMBRC platform PIQv for image analysis and CCPv for samples storage and supported by EMBRC-France, whose French state funds through the ANR within the Investments of the Future program under reference ANR-10-INBS-02. The Tara Pacific expedition would not have been possible without the participation and commitment of over 200 scientists, sailors, artists and citizens (see https://zenodo.org/record/3777760#.YfEEsfXMLjB). This publication is number X of the Tara Pacific Consortium.

## Author contributions

Conceptualization and methodology: FL, GB, SP, EB, NC, ED, JMF, SGJ, IK, MLP, JPMP GR, SR, ER, AV, CRV, BB, CB, DF, PF, PEG, EG, SR, SS, OT, RT, RVT, PW, DZ, DA, SP, MBS, CdV, EB, GGSample collection: FL, GB, SP, SA, EB, PC, ODS, ED, AE, JMF, JFG, BCH, YL, RM, DAP, MLP, JP, GR, SR, ER, CRV, GI, DF, PF, PEG, EG, SR, SS, OT RVT, PW, DZ, DA, SP, CdV, EB, GG

Samples analysis and data analysis: FL, GB, SA, EB, NC, MC, PC CD, ED, AE, JF, JMF, JFG, BCH, LJ, SGJ, RLK, YL, DM, RM, ZM, NM, DAP, MLP, MP, JR, GR, SR, ER, CRV, BB

Data production (models/satellites): FL, GB, AB, ODS, CRV

Data Curation and validation: FL, GB, SP, AB, MC, ODS, CD, ED, AE, JF, JMF, LJ, SGJ, RLK, IK, YL, DM, RM, ZM, NM, MP, JR, GR, ER, AV, CRV

Funding Acquisition: FL, NC, PC, ED, JF, JMF, JFG, SGJ, IK, DM, NM, MLP, MP, GR, ER, AV, CRV, BB, CB, DF, PF, PEG, EG, SR, SS, OT, RT, RVT, PW, DZ, DA, SP, CdV, EB, GG

Project Administration and supervision: FL, SP, SA, EB, JMF, ER, CRV, CM, BB, CB, DF, PF, PEG, EG, SR, SS, OT, RT, RVT, PW, DZ, DA, SP, MBS, CdV, EB, GG

Visualization: FL, GB, ZM

Writing – Original Draft Preparation: FL, GB, SP, SA, AB, EB, NC, MC, ODS, ED, JF, JMF, SGJ, RLK, RM, ZM, CRV

All authors have read and reviewed the manuscript.

